# NKG2A and HLA-E define a novel alternative immune checkpoint axis in bladder cancer

**DOI:** 10.1101/2022.03.04.482960

**Authors:** Bérengère Salomé, John P. Sfakianos, Jorge Daza, Andrew Charap, Christian Hammer, Romain Banchereau, Adam M. Farkas, Daniel Geanon, Geoffrey Kelly, Ronaldo M. de Real, Brian Lee, Kristin G. Beaumont, Sanjana Shroff, Yuan Shuo A. Wang, Ying-chih Wang, Tin Htwe Thin, Monica Garcia-Barros, Everardo Hegewisch-Solloa, Emily M. Mace, Li Wang, Timothy O’Donnell, Diego Chowell, Ruben Fernandez-Rodriguez, Mihaela Skobe, Nicole Taylor, Seunghee Kim-Schulze, Robert P. Sebra, Doug Palmer, Eleanor Clancy-Thompson, Scott Hammond, Alice O. Kamphorst, Karl-Johan Malmberg, Emanuela Marcenaro, Pedro Romero, Rachel Brody, Mathias Viard, Yuko Yuki, Maureen Martin, Mary Carrington, Reza Mehrazin, Peter Wiklund, Ira Mellman, Sanjeev Mariathasan, Jun Zhu, Matthew D. Galsky, Nina Bhardwaj, Amir Horowitz

## Abstract

PD-1/PD-L1-blockade immunotherapies have limited efficacy in the treatment of muscle-invasive bladder cancer (MIBC) and metastatic urothelial carcinoma. Here, we show that *KLRC1* (NKG2A) expression associates with improved survival and responsiveness to PD-L1 blockade immunotherapy in *CD8A*^high^ bladder tumors. The loss of antigen presentation is a common mechanism for tumor escape in bladder cancer. NKG2A^+^ CD8 T cells are able to circumvent HLA-ABC loss through TCR-independent cytotoxicity, which is partly mediated by DNAM-1. In bladder tumors, NKG2A is acquired on a subset of PD-1^+^ CD8 T cells, alongside stronger tissue-residency memory features, TCR-independent cytotoxicity and evidence of recent proliferation. HLA-E is low but variably expressed on bladder tumors. When expressed, NKG2A^+^ CD8 T cell anti-tumor responses to HLA-ABC-deficient tumors are inhibited and partly restored upon NKG2A blockade. Overall, our study identifies an alternative path for CD8 T cell exhaustion, that is mediated by NKG2A upregulation and TCR-independent cytotoxicity.

## Introduction

Resistance mechanisms are commonly employed by tumors in response to immune pressures exerted by effector cells. Among such resistance mechanisms, increased expression of inhibitory receptors on CD8^+^ T cells constrains their anti-tumor cytolytic potential (Thommen and Schumacher, 2018). Monoclonal antibodies that abrogate these inhibitory interactions have transformed the landscape for treatment of solid tumors (Bajorin et al., 2021; Chao et al., 2021; Finn et al., 2020; Motzer et al., 2020; Rittmeyer et al., 2017; Schmid et al., 2020; Topalian et al., 2019) and hematological malignancies (Chen, 2019). Immunotherapies targeting the PD-1/PD-L1 pathway have particularly proven effective in controlling tumor growth through the reinvigoration of CD8^+^ T cells across numerous tumor settings. PD-1-mediated inhibition not only targets TCR signaling but also CD28 (Boussiotis, 2016; Hui, 2017; Yokosuka et al., 2012), the latter being an important key player for the induction of effective anti-tumor CD8^+^ T cell responses following PD-1 blockade (Hashimoto, 2018; Kamphorst, 2017). Further, the loss of HLA class I expression on tumors is a common evasion strategy framing an important need to better understand CD28/TCR-independent mechanisms of anti-tumor functions.

Advanced urothelial cancer of the bladder is an aggressive malignancy with a median survival of approximately 13-16 months for patients with metastatic disease (Galsky et al., 2020). Several PD-1 and PD-L1 inhibitors have been approved by the US Food and Drug Administration (FDA) for bladder cancer treatment since 2016, spanning non-muscle-invasive to metastatic disease states. While responses to PD-1/PD-L1 blockade are often durable relative to other systemic cancer treatments, such responses are achieved in only 15-25% of patients with bladder cancer (Patel et al., 2020). The observed efficacy holds promise but suggests additional adaptive resistance mechanisms and provides strong rationale for targeting additional immune checkpoint axes in bladder cancer (Rosenberg et al., 2016). Several important intrinsic and extrinsic mechanisms of resistance to checkpoint blockade immunotherapies have been characterized (Fares, 2019), such as a tumor’s intrinsic ability to alter its recognition as ‘foreign’ by immune cells or rather extrinsic to the tumor microenvironment (TME), such as immunosuppressive cytokines produced by immune and stromal cells and alterations to vasculature and extracellular matrix. High tumor mutation burden (TMB) is more likely to generate neoantigens that can be recognized by adaptive cytotoxic T cells, and TMB correlates with improved treatment response across numerous tumor indications, including bladder cancer (Cristescu et al., 2018). However, our group recently demonstrated that an anti-tumor adaptive immune response may co-exist with a pro-tumorigenic inflammatory response, and that the balance between these two signatures (coined 2IR score) strongly stratifies responders and non-responders in metastatic bladder cancer patients treated with anti-PD-1/PD-L1 blockade (Wang et al., 2021).

HLA-E is a non-classical MHC class I molecule that is ubiquitously expressed on hematopoietic cells at low abundance and binds to the heterodimeric complex CD94/NKG2A (Boyington et al., 1999; Carretero et al., 1997; van Hall et al., 2019). While the CD94 c-type lectin-like receptor engages the majority of contact points with HLA-E, it relies on its structural binding partner, NKG2A, to mediate the inhibitory signal to NK cells and to CD8 T cells through two intracellular tyrosine-based inhibition motifs encoded on the cytoplasmic tail of NKG2A (Aramburu, 1990; Brooks, 1997; Speiser, 1999). A humanized monoclonal antibody binding to the NKG2A receptor, monalizumab, has been developed, and numerous clinical trials are ongoing across multiple tumor indications (NCT04590963, NCT02643550, NCT04307329, NCT02557516, NCT02671435, NCT04333914, NCT03822351, NCT04145193,

NCT03794544, NCT03833440, NCT03088059) (André et al., 2018; van Hall *et al.*, 2019). *In vitro* blockade of NKG2A, alone, or in combination with targeting the PD-1 pathway stimulates NK cell functions but is collectively required to stimulate a strong CD8 T cell response to HLA-E^+^ PD-L1^+^ tumors. Combined administration of anti-NKG2A and anti-PD-L1 blocking antibodies unleashes NK and CD8 T cells and subsequently slows tumor progression in mouse models and preliminary analyses suggest *in vivo* efficacy of monalizumab when in combination with the EGF receptor blockade antibody (cetuximab) in recurrent/metastatic head and neck squamous cell carcinoma (HNSCC) (André *et al.*, 2018).

In this study, we demonstrate that muscle-invasive bladder tumors have significantly reduced expression of MHC class I molecules, potentially limiting CD8 T cell responses to the tumor. We describe how a subset of NKG2A-expressing CD8 T cells can overcome this evasion strategy through TCR-independent functions that allow them to degranulate and produce critical soluble inflammatory mediators in response to MHC class I-deficient tumors, partly due to their increased expression of activating receptors, such as the DNAX Accessory Molecule (DNAM)-1/CD226 as well as due to loss of CD28 and reduced sensitivity to PD-1-mediated signaling. We further demonstrate that the NKG2A^+^ CD8 T cell ‘missing-self’ response is inhibited by HLA-E but restored by NKG2A blockade. HLA-E expression is strongly downregulated but variable on bladder tumors and highly significantly correlates with NKG2A blockade efficiency, providing a strong rationale for the use of NKG2A blockade for treatment of HLA-E^+^ bladder tumors. This study will additionally provide insights into engineering CD8 T cells for adoptive cell therapies.

## Results

### NKG2A expression associates with better survival in muscle-invasive bladder cancer

Based on the increased interest into NKG2A function in NK and CD8 T cells and the emerging data in clinical trials of NKG2A blockade in other tumor types, we first evaluated the effect of *KLRC1* (NKG2A) gene expression on the survival of cancer patients from The Cancer Genome Atlas (TCGA) “BLCA” (Bladder cancer), “KIRC” (Kidney clear cell carcinoma), “SKCM” (melanoma), “LUAD” (lung adenocarcinoma), “LUSC” (lung squamous cell carcinoma) and “OV” (ovarian cancer) cohorts. These cohorts are characterized by different tumor biology, different treatments, and different tumor stages. We stratified the cohorts into quantiles based on *KLRC1* expression and measured the effects of *KLRC1*^high^ (>75%) vs *KLRC1*^low^ (<25%). *KLRC1* expression significantly associated with better overall survival of the bladder cancer patients (Figure 1A). The TCGA bladder cancer database represents tumor specimens from 406 patients, most of them with muscle-invasive stage (n=371 MIBC, n=5 NMIBC, n=30 unknown pathology). We then divided the bladder cancer cohort based on the median expression of *CD8A* (CD8) and *PDCD1* (PD-1). *KLRC1* effect on survival is restricted to patients with high expression of *CD8A* or *PDCD1* in the tumor (Figure 1B-C). Additionally, we considered *NCR1* (NKp46) expression to better distinguish NK and CD8 T cells. However, the TCGA cohort size does not allow for evaluation of *KLRC1* effect on survival based on *NCR1* expression, as only eleven patients were found to display *NCR1*^low^ *KLRC1*^high^ expression (data not shown). *KLRC1* did not associate with patient survival in the KIRC (*KLRC1*high versus low: p=0.08), SKCM (p=0.1), LUAD (p=0.7), LUSC (p=0.7), OV (p=0.4) cohorts (data not shown). Overall, these results suggest a beneficial role for NKG2A on CD8 T cells in bladder cancer.

**Figure 1.**
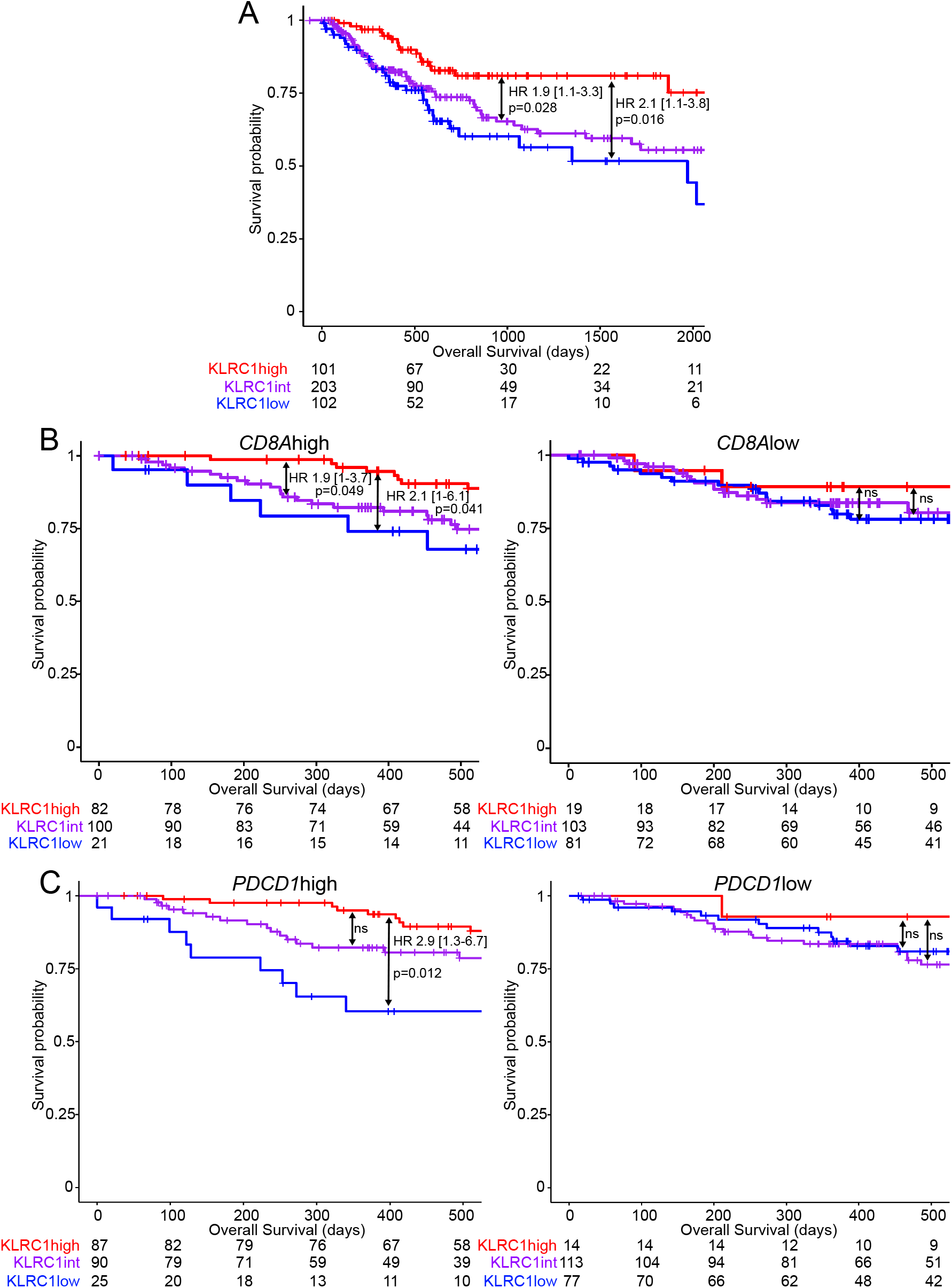
*KLRC1* (NKG2A) gene expression correlates with better overall survival in bladder cancer. Kaplan-meier survival curves of the TCGA bladder cancer cohort showing the association between *KLRC1* (NKG2A) expression and overall survival in (A) all patients, or patients with (B) *CD8A*^high^ or *CD8A*^low^ (CD8) or (C) *PDCD1*^high^ or *PDCD1*^low^ (PD-1) gene expression. Hazard ratios and p-values were calculated using COX proportional hazards regression models. These models were corrected for age and number of somatic mutations.

### Bladder tumors have reduced expression of HLA class I but retain expression of ligands for DNAM-1

To better understand the tumor microenvironment (TME) in bladder cancer, we performed single-cell RNA sequencing on fresh bladder tumors from 8 patients (3 non-muscle invasive NMIBC, 5 muscle-invasive MIBC) (Table 1). Using UMAP clustering analyses, we distinguished clusters of tumor cells, fibroblasts and major immune populations (Figure 2A, Table S1). To gain insights into antigen presentation capacity of bladder tumors to CD8 T cells, we compared the expression of genes encoding for HLA class I and beta-2 microglobulin (*B2M*) between the tumors and the immune cell clusters. We found that expression levels of *HLA-A*, *HLA-B*, *HLA-C*, *HLA-E*, *B2M* were significantly reduced on the tumor cells compared to the immune compartment in each of the 8 patients (Figure 2B). Flow cytometric analysis of HLA class I expression on CD45^+^ peripheral blood mononuclear cells (PBMCs) (n=19) and on CD45^−^ and CD45^+^ cells from tumors (n=25) of bladder cancer patients confirmed a significant reduction in cell-surface expression of HLA-ABC and HLA-E on the CD45^−^ tumor cells compared to the CD45^+^ immune compartment (Figure 2C). Additionally, HLA-E, but not HLA-ABC expression was reduced on intratumoral immune cells compared to their peripheral blood counterparts, suggesting distinct regulatory mechanisms governing the expression of these HLA molecules within the TME. Muscle-invasive specimens displayed significantly lower expression of HLA-ABC on tumor cells compared to non-muscle-invasive specimens (Figure 2D). A similar trend was observed for HLA-E, although no statistical significance was reached.

**Table 1.**
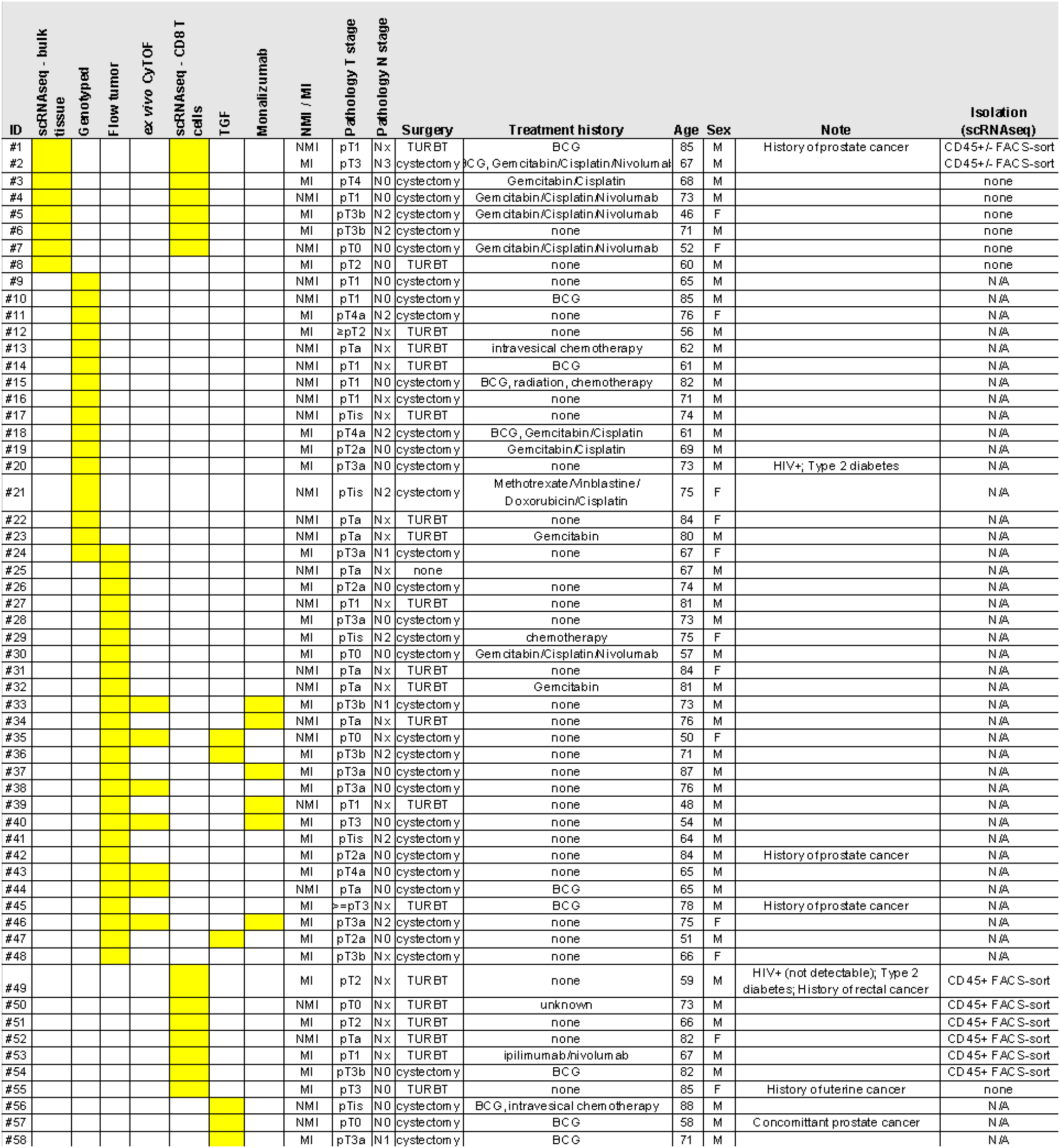
Patient information

**Figure 2.**
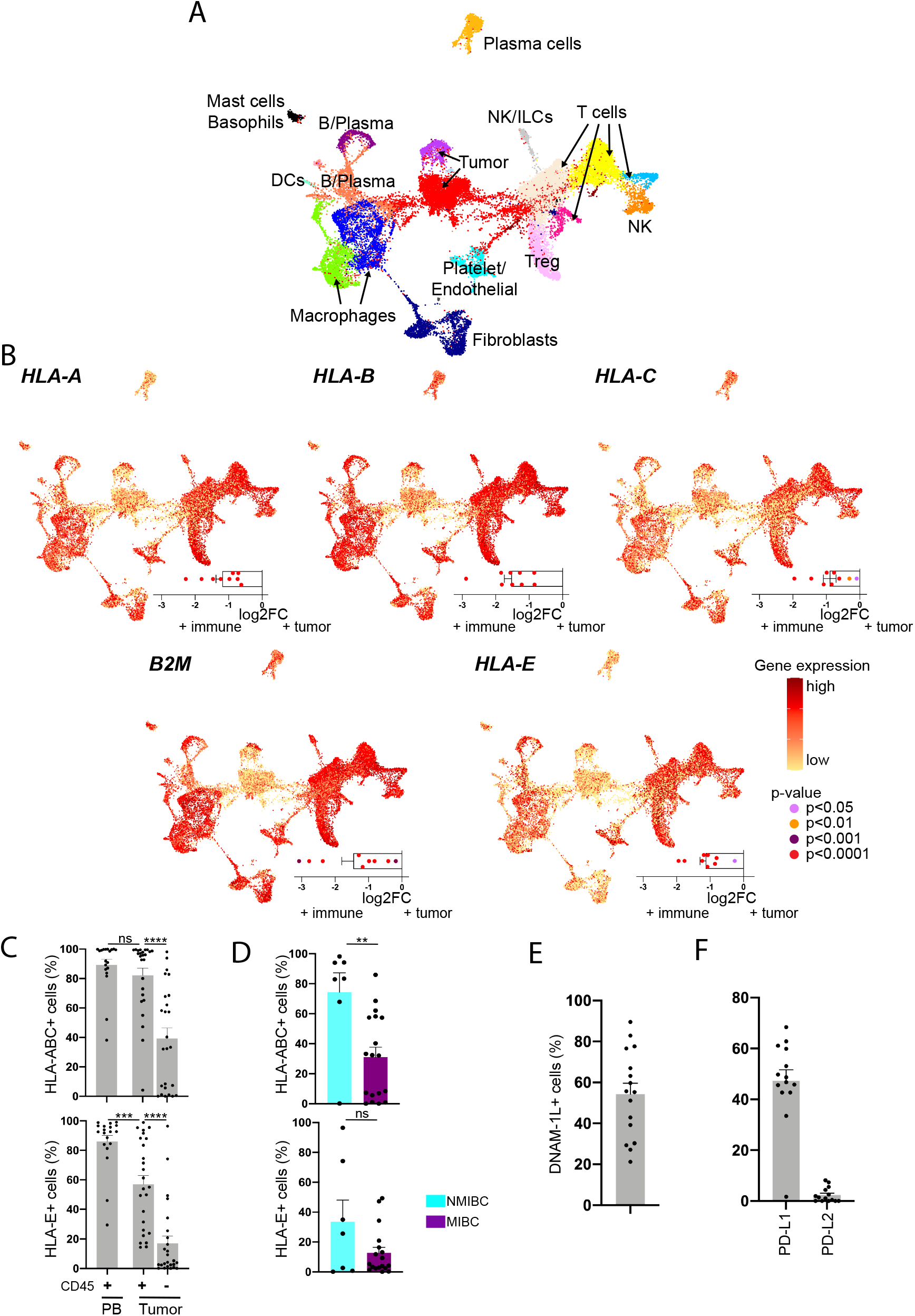
Bladder tumors have reduced expression of HLA class I but retain expression of ligands for DNAM-1. (A-B) Single-cell RNA sequencing data was generated from tumors of bladder cancer patients (n=8). (A) UMAP clustering analysis. Each color represents a cluster. (B) For each gene, the central graph shows the gene expression level on the UMAP clusters while the bottom right graph shows the log2(Fold change) of the gene expression between the tumor cells and the immune cells. (C-F) Flow cytometry analyses were performed on PBMCs and tumor-dissociated cells of bladder cancer patients. (C) Frequency of cells expressing HLA-ABC or HLA-E among CD45^+^ PBMCs (n=19) and CD45^+^/CD45^−^ cells from the tumor site (n=25). (D) Frequency of CD45- tumor cells expressing HLA-ABC or HLA-E in NMIBC (n=7) and MIBC (n=18) patients. (E) Frequency of CD45- tumor cells expressing at least one DNAM-1 ligand (CD112 and/or CD155) (n=16). (F) Frequency of CD45^−^ tumor cells expressing PD-L1 or PD-L2 (n=14). Paired t-tests were used in (C) and unpaired t-test in (D). “ns” p>0.05, ** p<0.01, *** p<0.001, **** p<0.0001.

Loss of HLA class I on tumors (loss of heterozygosity (LOH)) is a common phenotype and can be driven by somatic mutations (Grasso et al., 2018) or by post-transcriptional or epigenetic changes in response to environmental pressures (Ritter et al., 2017). Loss of HLA-A, −B or −C allotypes promotes a mechanism of tumor evasion from anti-tumor CD8 T cells (Garrido et al., 2016). We performed an allele-specific copy number FACETS analysis (Shen and Seshan, 2016) of the TCGA “BLCA” cohort and observed 21.8% of patients had lost one or more alleles of HLA class I. Further, we genotyped HLA class I alleles in patient-matched germline and tumor tissues from our cohort of bladder cancer patients and did not observe LOH in any of the tumor or urine specimens compared to germline tissue (Table S2) suggesting that somatic mutations in the *HLA class I* locus are not the dominant driver for the observed loss in expression of HLA class I on bladder tumors.

We further characterized an historical panel of bladder tumor cell lines by flow cytometric analysis for expression of activating and inhibitory ligands that are conventionally ascribed to modulating NK and T cell functions. The panel of tumor lines represents a broad spectrum of clinical grades (G1-G3), as established according to the 1973 WHO classification (Mostofi, 1973). We found NKp44- and NKp46-ligands (L) to be absent from all 11 bladder cancer lines and NKp30-ligands to be expressed only at low levels on the RT-4 tumor cell line (Figure S1). Conversely, all cell lines expressed at least one NKG2D-L among ULBP1, ULBP2/5/6, ULBP3, MICA, MICB and had high expression of DNAM-1 ligands (CD112, CD155). We also noted expression of DNAM-1 ligands on primary bladder tumors (Figure 2E). Finally, whereas PD-L1 and PD-L2 were expressed on the bladder tumor cell lines, we observed PD-L1, but not PD-L2, on primary tumors (Figure 2F). Collectively, these results suggest that even though the bladder TME might be impaired in pathways for antigen presentation to CD8 T cells, other pathways known to initiate NK cell “missing-self” reactivity are present.

### NKG2A-expressing CD8 T cells are differentiated and possess TCR-independent anti-tumor functions

To better understand the phenotype of CD8 T cells that may be engaged independent of MHC class I, we sought to characterize NKG2A^+^ CD8 T cells. We evaluated the *ex vivo* phenotypes and functions of NKG2A^+^ CD8 T cells initially in the blood of healthy individuals (n=20) using mass cytometry (Table S3, panels 1-3). We defined CD8 T cells as CD45^+^ CD4^−^ CD14^−^ CD19^−^ Vα24/Jα28TCR^−^ γδTCR^−^ CD3^+^ CD8^+^, thereby excluding invariant NKT cells and γδT cells. We distinguished naïve cells as CD45RA^+^ CCR7^+^, effector memory (EM) cells as CD45RA^−^ CCR7^−^, central memory (CM) cells as CD45RA^−^ CCR7^+^, and EMRA cells as CD45RA^+^ CCR7^−^, We observed NKG2A to be restricted to the memory CD8 T cell compartment, with highest expression at the effector (EM) and central memory (CM) stages (Figure 3A). NKG2A expression did not depend on gender, age or prior infection with cytomegalovirus (Figure S2A-B). Further, while cell-surface expression of HLA-E on peripheral blood mononuclear cells (PBMCs) varies considerably in healthy individuals and influences the expression of NKG2A on NK cells as well as NKG2A^+^ NK cell function (Horowitz et al., 2016; Ramsuran et al., 2018), we observed that NKG2A expression on CD8 T cells did not depend on HLA-E expression (Figure S2C). We then compared the *ex vivo* phenotypes of NKG2A^+^ and NKG2A^−^ CD8 T cells at each memory stage. NKG2A^+^ CD8 T cells displayed higher levels of the NK-activating receptor DNAM-1 and the canonical lineage marker CD56 (Figure 3B). DNAM-1 and NKG2D were the most highly expressed NK receptors on NKG2A^+^ CD8T cells, though only DNAM-1 expression differed between NKG2A^+^ and NKG2A^−^ CD8 T cells (Figure 3B, S2D). TIGIT and PD-1 were expressed at lower levels on NKG2A^+^ CD8 T cells at the EM and CM stages. At the EMRA and CM stages, NKG2A^+^ CD8 T cells also expressed the highest levels of the transcription factor (TF) T-bet. However, unique to the EMRA stage, NKG2A^+^ CD8 T cells expressed significantly higher levels of eomes as well as a significantly higher cytolytic payload (e.g. Granzyme A, Granzyme B, Perforin).

**Figure 3.**
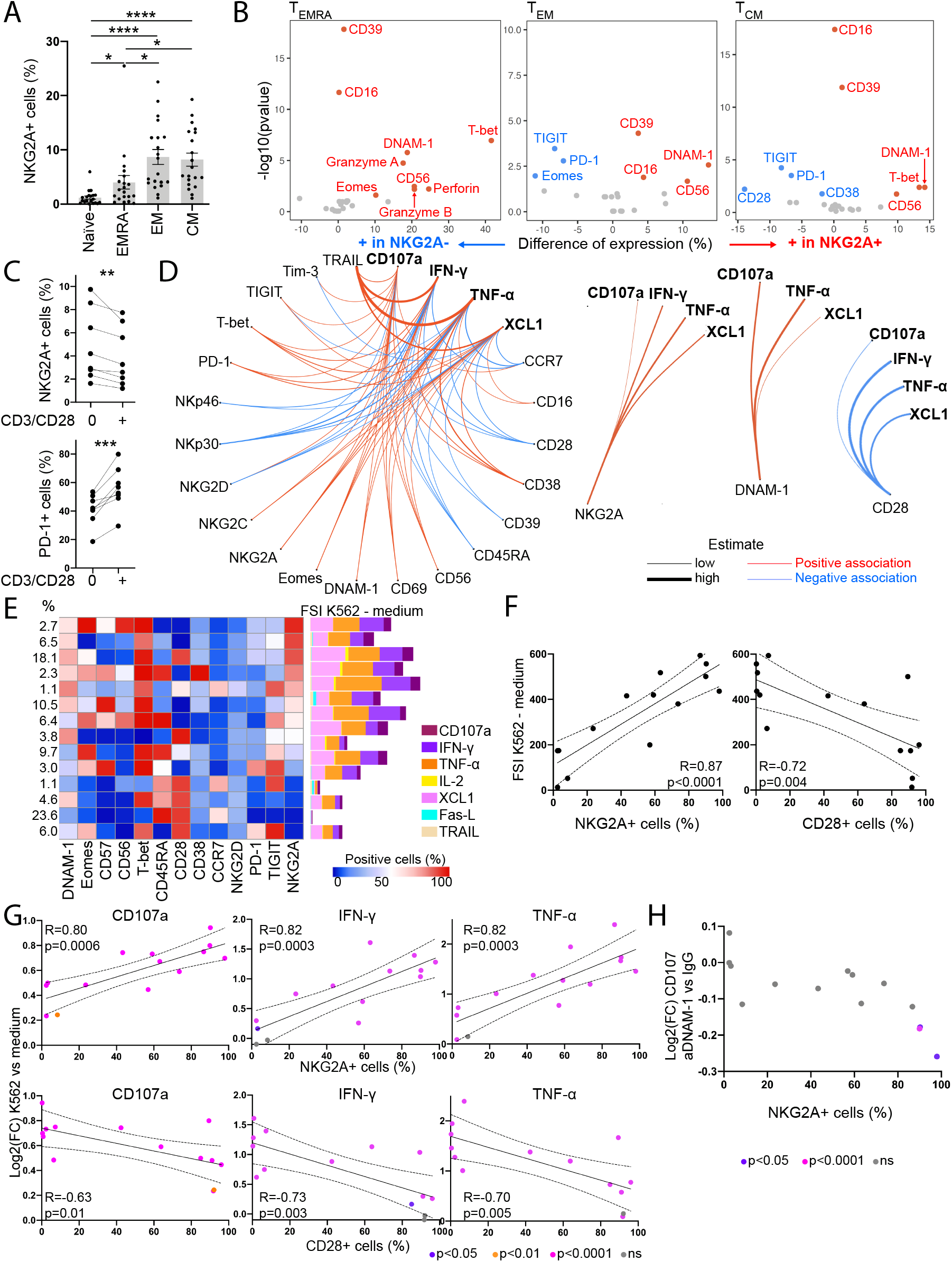
NKG2A-expressing CD8 T cells are differentiated and possess TCR-independent NK-like functions in healthy individuals. (A-B) Mass cytometry was performed *ex vivo* on HD PBMCs (n=20). (A) NKG2A expression on CD8T cells from HD PBMCs stratified according to the stage of CD8 T cell differentiation. (B) Differentially expressed markers between NKG2A^+^ and NKG2A^−^ CD8 T cells in HD PBMCs. (C) NKG2A and PD-1 expression on CD45RA^−^ CD8 T cells after 24h culture of CD8 T cells from HD PBMCs with low-dose IL-2, IL-7, IL-15, with or without CD3/CD28 beads. (D) CD8 T cells from HD PBMCs were recovered overnight with low-dose IL-12, IL-15, IL-18 prior to a 5h co-culture with K562 (n=20). Association between expression of phenotypic and functional markers following co-culture with K562 tumors. (E-H) CD8 T cells from HD PBMCs were recovered overnight with low-dose IL-12, IL-15, IL-18 prior to a 5h co-culture with K562 in the presence or absence of anti-DNAM-1 blocking antibody (n=10). (E) FlowSOM cluster analysis of CD8 T cells. The heatmap displays 14 clusters (rows) expressing various combinations of markers (columns). The distribution of CD8 T cells among these clusters is represented at the left. The Functional Strength index (FSI) upon K562 stimulation minus the FSI upon 5h culture in R10 medium is displayed at the right of the heatmap. (F) Correlation between the frequency of NKG2A^+^ or CD28^+^ CD8 T cells within each cluster and FSI upon K562 stimulation minus FSI in the absence of K562. (G) Correlation between the frequency of NKG2A^+^ or CD28^+^ CD8 T cells within each cluster and Log2(Fold change) of the expression of CD107a, IFN-g, and TNF-a upon K562 stimulation. (H) Log2(Fold change) of CD107a expression upon addition of anti-DNAM-1 blocking antibody, compared to isotype control antibody, in all clusters. Paired t-tests were used in (A), (B), (C), linear models in (D), Pearson correlations in (F), (G) and unpaired t-tests in (G), (H). All statistical tests were adjusted for multiple comparison. Only the significant comparisons are displayed in (D). * p<0.05, **** p<0.0001.

Strikingly, *ex vivo* expression of CD28 was downregulated on CM CD8 T cells when NKG2A was expressed (Figure 3B, S2E). To better understand the relationship between CD28 and NKG2A, we cultured PBMCs for 24 hours in the presence of low dose cytokines and CD3/CD28 stimulation. We observed decreased expression of NKG2A with a concomitant increase in PD-1 expression on CD8 T cells, suggesting that the CD28 and NKG2A pathways are mutually exclusive (Figure 3C). We next assessed the CD8 T cell response to HLA class I-deficient K562 tumor cells. CD8 T cells were isolated from blood of healthy individuals (n=20), cultured overnight in low-dose type 1 cytokines and co-cultured for 5h with K562. CD8 T cells degranulated and produced TNF-α, IFN-γ, XCL1 and IL-2 in response to K562 (Figure S2F). Although degranulation was observed at all differentiation stages, efficient release of cytotoxic mediators (Granzyme A, Granzyme B, Perforin) was only observed at the EMRA stage (Figure S2F-G). Using linear models, we characterized associations between CD8 T cell phenotypes and reactivity to K562 (Figure 3D). Specifically, the response to K562 was restricted by CD28 while enhanced by DNAM-1 and NKG2A.

Additionally, we observed that the association between NKG2A expression and TCR-independent NK-like functions in CD8 T cells is tightly regulated by HLA-E, being restricted in functional capacity in healthy individuals with endogenously higher cell-surface expression of HLA-E (Figure S2H). Thus, the data suggest that while HLA-E expression does not affect the numbers of circulating NKG2A^+^ CD8 T cells, higher expression of HLA-E strengthens their inhibition.

In order to identify potential mechanisms underlying TCR/CD28-independent CD8 T cell functions, we performed a cluster analysis of CD8 T cells from healthy individuals (n=10) co-cultured with K562 tumors in the presence or absence of blocking antibodies targeting activating NK cell receptors (Figure 3E). We identified 14 relevant clusters, including 3 clusters with >95% of NKG2A^+^ cells and 4 clusters with <2% NKG2A^+^ cells. The majority of NKG2A^+^ CD8 T cell clusters expressed the highest levels of DNAM-1. We then assessed the CD8 T cell response to K562 tumors in each of these clusters by calculating a functional stimulation index (FSI), which considers the frequency and signal intensity of each functional marker. An increasing FSI acquired upon K562 co-culture reflects a CD8 T cell polyfunctional response to HLA-ABC-deficient tumors. CD8 T cell polyfunctionality positively correlated with NKG2A expression and negatively correlated with CD28 expression within each cluster (Figure 3F). The functional pathways affected by NKG2A and CD28 expression are, in particular, TCR-independent degranulation and production of TNF-α and IFN-γ (Figure 3G, S2I).

Blocking antibodies targeting the NK-activating receptors NKp30, NKp46, NKG2D or DNAM-1 were next used to treat CD8 T cells before co-culture with K562 tumors. Abrogating the interactions of NKp30, NKp46 or NKG2D with their activating ligands had no effect on the FSI (data not shown). However, blocking DNAM-1 strongly impaired CD8 T cell TCR-independent function, with its effects being restricted to CD8 T cell degranulation (Figure 3H, S2J). Strikingly, the effects of DNAM-1-blockade were confined to the three clusters with the highest frequency of NKG2A^+^ CD8 T cells. A linear mixed model (NKG2A*anti-DNAM-1 effect: p=0.010) was performed, which confirmed that the effects of DNAM-1-blockade on CD8 T cell degranulation are contingent upon expression of NKG2A. Collectively, the data indicate that NKG2A marks a program of TCR-independent, DNAM-1-mediated “missing-self” degranulation by CD8 T cells in the blood of healthy individuals.

### NKG2A defines an alternatively exhausted subset of TRM CD8 T cells that retain TCR-independent anti-tumor functions in bladder tumors

In order to confirm the role of NKG2A in bladder cancer, we next sought to evaluate NKG2A acquisition on CD8 T cells in bladder tumors using mass cytometry (Table S3, panel 4). We analyzed CD8 T cell phenotypes in matched samples of bladder draining lymph nodes (BDLN) (n=5), bladder tumors (n=7) and adjacent, non-involved bladder tissue (n=6) (Table 1). Hierarchical clustering analyses revealed CD8 T cell clusters that were defined by their variation in expression of markers associated with tissue residence (CD49a, CD69, CD103), differentiation (Eomes, T-bet, Tox, TCF-1), effector functions (granzymes A and B, perforin) and exhaustion (CD39, NKG2A, PD-1, TIGIT, Tim-3) (Figures 4A-4C, and S3A). We observed a strong downregulation of CD28 expression concomitant to an upregulation of expression of tissue-residence markers, checkpoint inhibitors (PD-1, CD39) and exhaustion-related TFs (TOX, T-bet) on CD8 T cells upon entry in the tumor. Of note, tumor infiltration of bladder-draining lymph nodes did not seem to impact CD8 T cell phenotypes (Figure S3B).

**Figure 4.**
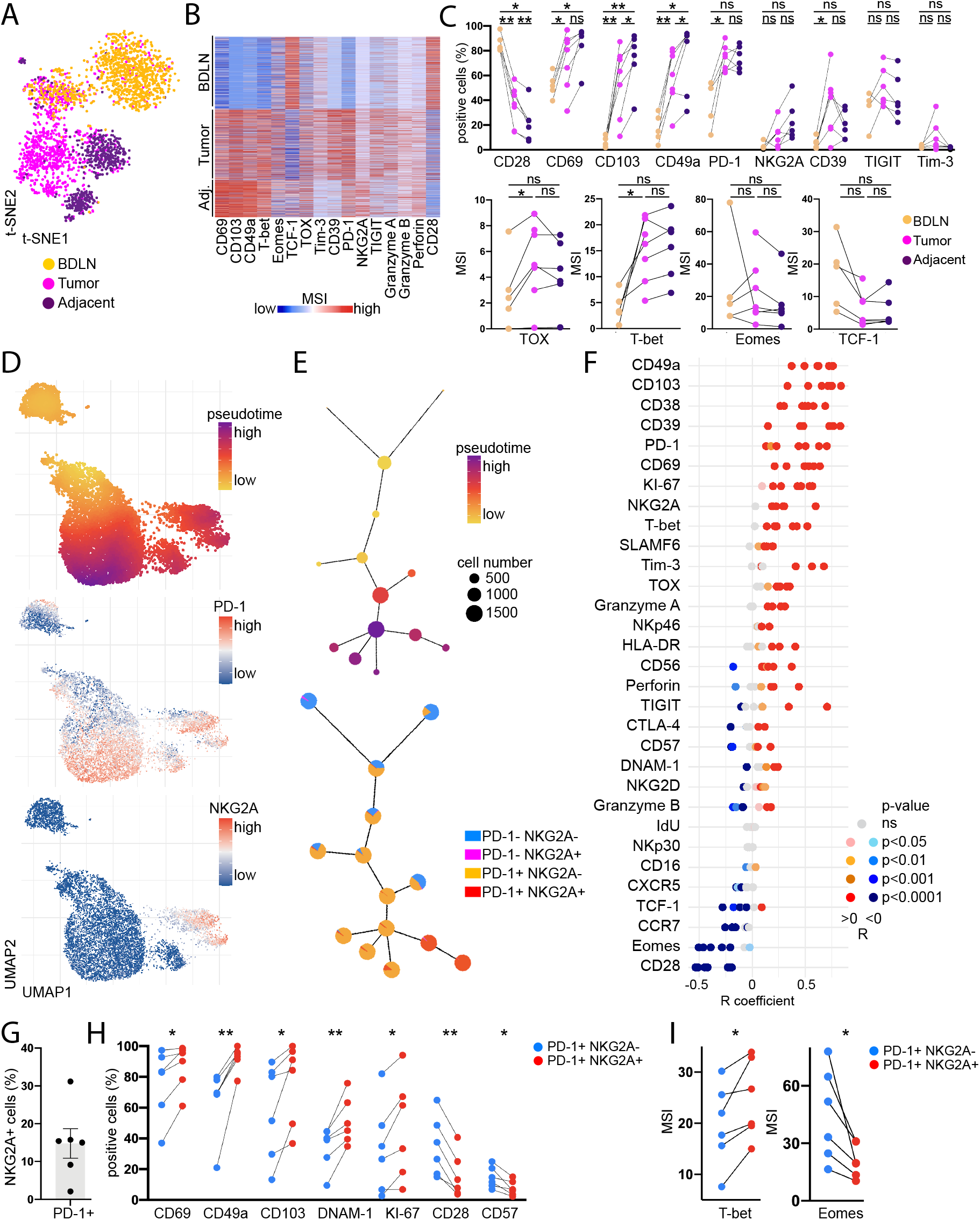
NKG2A defines an alternatively exhausted subset of TRM CD8^+^ T cells that retain TCR-independent functions in bladder tumors. (A-I) Mass cytometry was performed *ex vivo* on bladder tumor-draining lymph nodes (BDLN), tumor and adjacent non-involved tissue from bladder cancer patients. (A) Representative t-SNE clustering analysis of CD8 T cells (n=1 patient). (B) Representative heatmap of CD8 T cell phenotype (n=1 patient). (C) CD8 T cell phenotype in bladder tumor-draining lymph nodes (n=5), tumor (n=7) and adjacent tissue (“Adj.”) (n=6). Lines show matching samples from a same donor. (D) Representative UMAP analysis of CD8 T cells from the tumor tissue (n=1). (E) Representative graph of the PD1/NKG2A cell expression on the CD8 T cells per UMAP cluster alongside the pseudotime trajectory (n=1). (F) Correlation of marker expression with pseudotime (n=7) (G) NKG2A expression on PD-1^+^ CD8 T cells (n=6) (H) Frequency of expression of markers in NKG2A^−^ PD-1^+/−^ CD8 T cells in the tumors (n=6). Only the significant markers are displayed. (I) Median Signal Intensity (MSI) of transcription factors in NKG2A^−^ PD-1^+/−^ CD8 T cells in the tumors (n=6). Only the significant markers are displayed. Paired t-tests were used in (C), (H), (I) and Spearman correlations in (F). All p-values were adjusted for multiple comparison. “ns” p>0.05, * p<0.05, ** p<0.01.

In order to better understand the phenotypic diversity within the repertoire of tumor-infiltrating CD8 T cells, we performed a pseudotime analysis of seven bladder tumors. Pseudotime analysis predicts the phenotypic relatedness and assigns a trajectory for visualization (Dai et al., 2021). CD8 T cells acquired PD-1 in a progressive manner within the pseudotime trajectory, while NKG2A acquisition defined a branch point directly derived from a subset of PD-1^+^ CD8 T cells, suggesting that PD-1 is acquired before NKG2A during chronic activation (Figures 4D and 4E). Expression of immune checkpoints (PD-1, NKG2A, CD39, Tim-3), tissue-residence markers (CD49a, CD103, CD69), TFs (TOX, T-bet) and the proliferation marker KI-67 additionally positively correlated with progression through the pseudotime trajectory (Figures 4F and S3C). Conversely, CD28 and Eomes expression were negatively correlated with the pseudotime trajectory.

NKG2A was expressed on 15% [2-31%] of PD-1^+^ CD8 T cells in bladder tumors (Figure 4G). Further, when gated on PD-1^+^ CD8 T cells, NKG2A expression is strongly correlated with enhanced proliferation and tissue-resident memory (TRM) features (Figure 4H). Importantly, NKG2A^+^ PD-1^+^ CD8 T cells also displayed higher expression of DNAM-1 and lower expression of CD28, in line with our observations in healthy individuals. T-bet was increased and Eomes was decreased on NKG2A^+^ PD-1^+^ CD8 T cells (Figure 4H and 4I). Collectively, the data suggest that NKG2A marks an alternative exhaustion path in CD8 T cells from bladder tumors, that is defined by a progressive loss of TCR co-stimulation, recent proliferation and tissue-residency.

### NKG2A expression is evenly distributed across the CD8 T cell clonal repertoire

Considering the enhanced recent proliferative activity observed in NKG2A-expressing PD-1^+^ CD8 T cells, we compared the clonality of *KLRC1*^low^ and *KLRC1*^high^ CD8 T cells using publicly available matched TCR-sequencing and single-cell RNA sequencing data on bladder tumors (n=7) and non-involved adjacent tissue (n=2) (Oh et al., 2020). We did not observe any significant differences in the clonalities between *KLRC1*^low^ and *KLRC1*^high^ CD8 T cells, with similar frequencies of cells expressing distinct TCRs (Figures S4A, S4B, and S4C). We then compared the TCRs expressed by at least two cells within the *KLRC1*^low^ and *KLRC1*^high^ groups. We centered this analysis on bladder tumor samples for which at least two TCRs are expressed in at least two *KLRC1*^high^ cells, excluding three tumor specimens in which only one TCR was found to be expressed in at least two *KLRC1*^high^ cells. 74% (67-83%) of the TCRs found in at least two *KLRC1*^high^ cells were also found in at least two *KLRC1*^low^ cells, showing very similar clonal expansion independent of NKG2A expression at the *mRNA* level (Figure S4D). Collectively, the data indicate that NKG2A expression on CD8 T cells does not associate with a skewed clonal repertoire.

### CD8 T cells with higher *KLRC1* (NKG2A) expression display stronger cytolytic potential at the earlier exhaustion trajectory in bladder tumors

We next characterized the effects of NKG2A expression on CD8 T cells at their transcriptomic level using single-cell RNA sequencing data from 14 fresh bladder tumors (Table 1). We identified 8 CD8 T cell clusters (Figure 5A), which distribution was independent of the tumor stage (Figure S5A). The “TGATA2” cluster was defined by high *GATA2* expression, the “Tc17” cluster by high *IL17A*, *RORA* and low *IFNG*, *EOMES*, *GATA3*, *TBX21*, consistent with the Tc17 population (Hamada et al., 2009; Huber et al., 2009) and the “T_PROLIF_” cluster by high *MKI67* transcript levels (Figure 5A-B). The “T_0_” cluster represented the least differentiated state and neighbored the *PDCD1*^low^ “T_HSP_” clusters, which were characterized by high expression of stress response genes, such as heat shock proteins, in accordance with metabolic activity in response to CD8 T cell trafficking into an immunosuppressive tumor micro-environment (Figure 5B-C). One of the two T_HSP_ clusters (“T_HSP_LYTIC_”) displayed higher expression of cytotoxicity-related genes. The two remaining clusters displayed intermediate *PDCD1* and *KLRC1* or high *PDCD1* and *KLRC1* expression; thereby, representing earlier and later exhaustion stages (“T_EARLY_”, “T_LATE_”).

**Figure 5.**
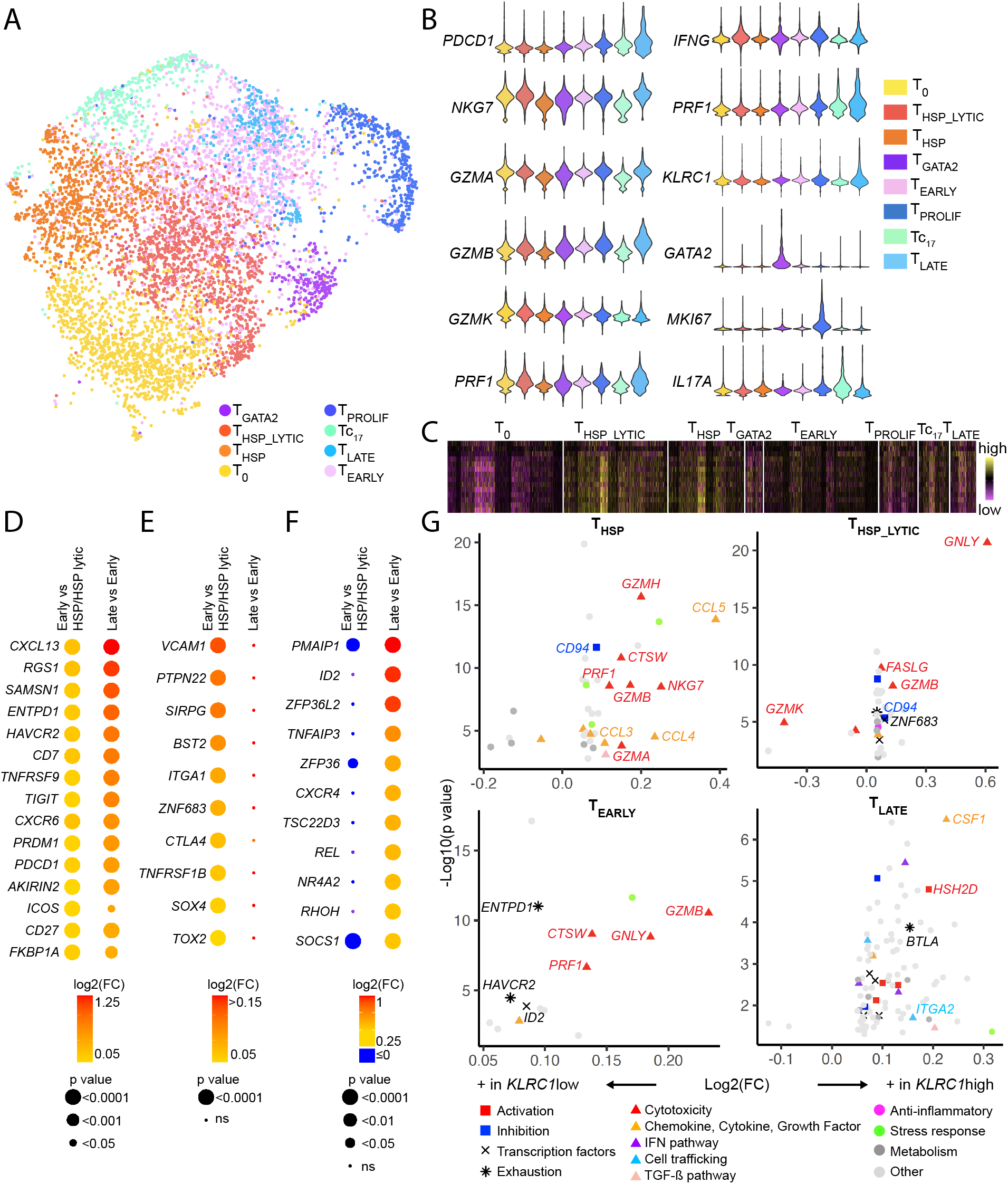
CD8 T cells with higher KLRC1 expression display stronger cytolytic potential at the earlier exhaustion stages in bladder tumors. (A-G) Single-cell RNA sequencing was performed on *ex vivo* bladder tumors (n=14). (A) UMAP clustering analysis of CD8 T cell gene expression. Each color represents one cluster. (B) Violin plots of gene expression per cluster. (C) Gene expression of upregulated stress-related genes in T_HSP_LYTIC_ and T_HSP_ versus T0 clusters. Genes displayed are (top to bottom order): *BAG1*, *GADD45G*, *DNAJB4*, *HSPA8*, *HSPE1*, *PPP1R15A*, *HSP90AB1*, *HSP90AA1*, *GADD45B*, *HSPH1*, *DNAJB1*, *KLF6*, *HSPA1A*, *HSPA1B*. Differential gene expression was performed on the T_EARLY_ versus T_HSP/HSP-LYTIC_ clusters and on the T_LATE_ versus T_EARLY_ clusters. A gene is considered upregulated if log2(FC)>0.05 and p<0.05. (D) Dot plots of genes that are upregulated in the T_EARLY_ versus T_HSP/HSP-LYTIC_ and TLATE versus T_EARLY_ clusters. (E) Dot plots of genes that are upregulated only in the T_EARLY_ versus T_HSP/HSP-LYTIC_ clusters. (F) Dot plots of genes that are upregulated only in the T_LATE_ versus T_EARLY_ clusters. (G) Volcano plots displaying differential expression of genes between KLRC1^high^ and KLRC1^low^ cells in the T_EARLY_,T_HSP_, T_HSP-LYTIC_ and T_LATE_ clusters.

We then evaluated the CD8 T cell transcriptome across the clusters, ranging from *PDCD1*^low^ T_HSP_/T_HSP_LYTIC_ stages to the *PDCD1*^int^ T_EARLY_ stage, as well as from the *PDCD1*^int^ T_EARLY_ to the *PDCD1*^high^ T_LATE_ stage. Non-relevant, non-coding or metabolism genes were excluded from this analysis. Fifteen genes were significantly upregulated, both, at the early (“T_EARLY_” vs “T_HSP_” and “T_EARLY_” vs “T_HSP_LYTIC_”) and late exhaustion stages (“T_EARLY_” vs “T_LATE_”), thereby capturing a range in CD8 T cell exhaustion (Figure 5D, Table S4). The defining genes included, but were not limited to, immune checkpoints (*ENTPD1, PDCD1, TIGIT, HAVCR2*), molecules involved in T cell migration (*CXCR6, RGS1*) and the TF Blimp-1 (*PRDM1*), which is associated with tissue-resident memory and exhaustion programs (Mackay, 2016). The early exhaustion stages (“T_EARLY_” vs “T_HSP_” and “T_EARLY_” vs “T_HSP_LYTIC_”) were distinguished by ten genes, including *CTLA4*, the exhaustion-related TF, *TOX2*, and tissue-residence-related genes (*ITGA1*, *ZNF683*) (Figure 5E) (Mackay, 2016; Seo et al., 2019). Unique to the late exhaustion stage (“T_LATE_” vs “T_EARLY_”) (Figure 5F, Table S4), we observed 126 upregulated genes, including genes involved in inhibition of T cell activation (*ZFP36, ZFP36L2, NR4A2, SOCS1)* or proliferation (*RHOH*) or in T cell apoptosis (*PMAIP1*), as well as the TF, *ID2*. These data confirm the association between tissue-resident memory and CD8 T cell exhaustion in bladder tumors and provide novel insights into modulation of critical CD8 T cell functions.

We next compared the transcriptome of *KLRC1*^high^ and *KLRC1*^low^ CD8 T cells in each of the four clusters. Only at the earliest exhaustion stages (“T_HSP_”, “T_HSP_LYTIC_”, “T_EARLY_”), *KLRC1*^high^ CD8 T cells contain higher transcript levels of genes encoding cytolytic mediators (*GZMA, GZMB, GZMH, GZMK, PRF1, CTSW, NKG7, GNLY*) (Figure 5G, Table S5). This association was lost by late-stage exhaustion (“T_LATE_”), where *KLRC1*^high^ CD8 T cells display, instead, higher levels of the inhibitory receptor *BTLA* and a gene inhibiting TCR/CD28 signaling (*HSH2D*) (Shapiro et al., 2004). *HSH2D* encodes the hematopoietic SH2 domain containing adaptor protein, also known as ALX, which inhibits transcriptional activation of the CD28 response element (RE) located within the IL-2 promoter during T cell activation. Collectively, these data provide evidence that early-exhaustion-pointed NKG2A-expressing CD8 T cells in bladder tumors possess a stronger capacity for cytolytic function while progressively suppressing TCR/CD28-mediated functions. Additionally, the data may suggest an overall shift by NKG2A-expressing CD8 T cells in prioritizing NK-like functions as tumors progressively lose HLA class I on their cell-surface.

### NKG2A acquisition on CD8 T cells defines a highly proliferative tissue-resident memory subset with enhanced response to HLA class I-deficient tumors

We next evaluated the effect of NKG2A and PD-1 acquisition on CD8 T cell phenotype and TCR-independent function in bladder tumors. In bladder tumors, TGF-β1 is produced and correlates with pathogenesis and aggressiveness of disease after surgery (Kim JH, 2001). TGF-β1 also induces PD-1 and NKG2A on CD3/CD28 -activated CD8 T cells in healthy donor peripheral blood (Gunturi et al., 2005; Park et al., 2016; Stefania Bertone, 2006). CD8 T cells from bladder tumor-draining lymph nodes are predominantly PD-1^−^ NKG2A^−^. In order to recapitulate PD-1 and NKG2A acquisition that occurs upon entry in the bladder tumors, we isolated CD8 T cells from bladder tumor-draining lymph nodes and expanded them for 8-13 days with CD3/CD28 tetramers and low-dose cytokines. We then isolated NKG2A^−^ PD-1^−^ CD69^−^ CD103^−^ CD49a^+/−^ CD8 T cells and expanded them for another three days with or without TGF-β1. TGF-β1 induced the acquisition of NKG2A and PD-1 (Figures 6A, S6A and S6B). We further co-cultured expanded CD8 T cells with HLA-deficient K562 tumors, resulting in degranulation and production of IFN-γ, TNF-α, XCL1 and IL-2 with no observed differences between CD49a^+^ and CD49a^−^ FACS-sorted cells (Figure 6B and S6C).

**Figure 6.**
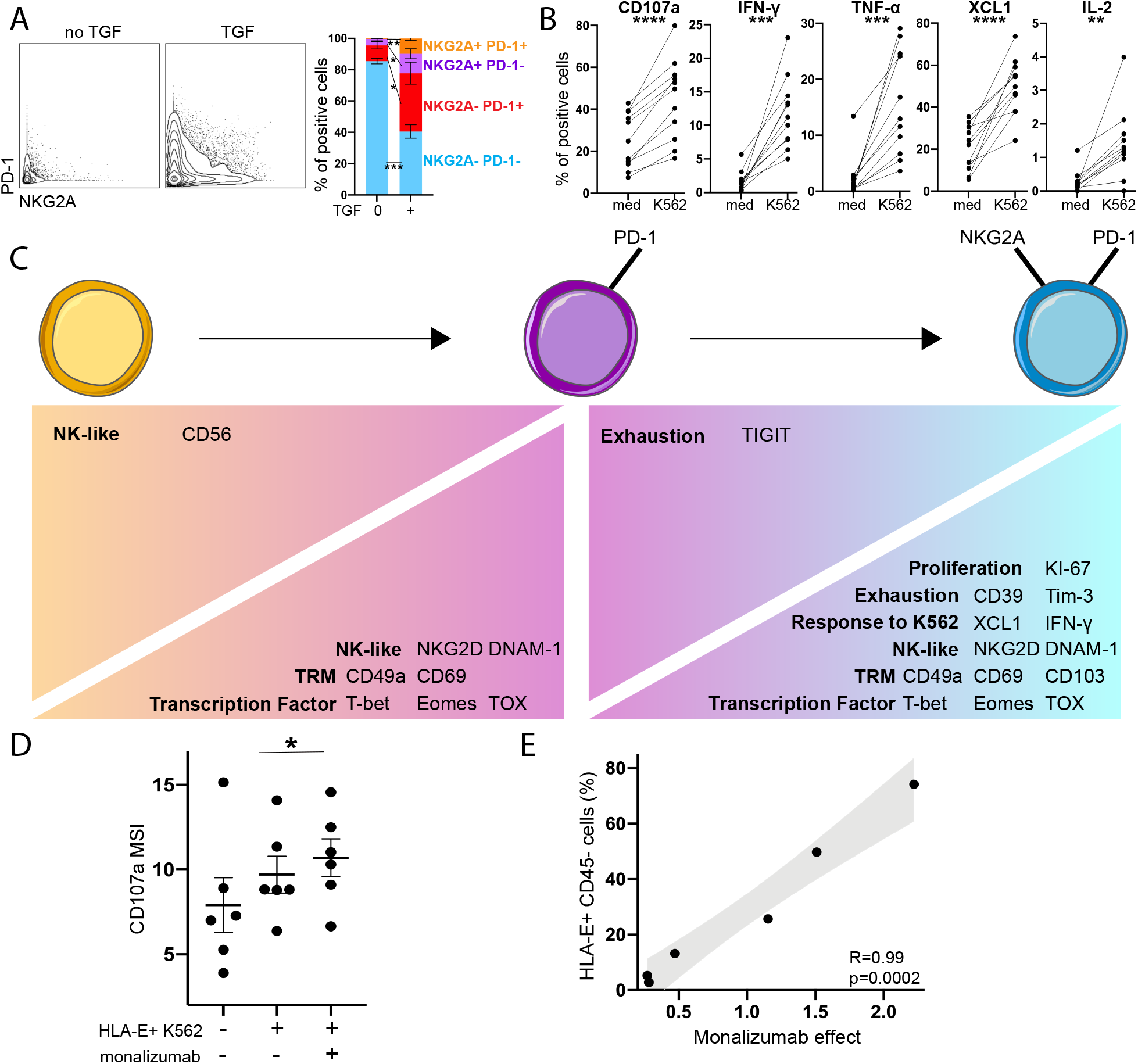
TCR-independent function is acquired by CD8 T cells upon NKG2A acquisition and is enhanced with NKG2A blockade in bladder cancer. (A-C) CD8 T cells were isolated from bladder tumor-draining lymph nodes and expanded for 8-13 days with IL-2, IL-7, IL-15 and CD3/CD28 tetramer. NKG2A^−^ PD1^−^ cells were then FACS-sorted and expanded during 3 additional days with or without TGF-β, prior to co-culture with K562. CyTOF was performed at all timepoints. (A) NKG2A/PD1 expression on CD49a^−^ NKG2A^−^ PD1^−^ CD8 T cells after 3-day culture with or without TGF-β (n=5). (B) Upregulation of functional markers upon K562 stimulation of CD49a^−^ and CD49a^+^ NKG2A^−^ PD1^−^ CD8 T cells after 3-day expansion in the presence of TGF-β (n=5 and 6, respectively). (C) Phenotype and response to K562 following 3-day expansion of NKG2A^−^ PD1^−^ CD8 T cells and NKG2A/PD1 upregulation (n=11). (D-E) CD8 T cells were isolated from bladder tumors (n=6) and expanded for 11-17 days with IL-2, IL-7, IL-15 and CD3/CD28 tetramer, prior to co-culture with HLA-E^+^ K562. (D) Effect of NKG2A blockade (monalizumab) on CD8 T cell degranulation upon HLA-E^+^ K562 stimulation. (E) Spearman correlation between HLA-E expression on CD45^−^ bladder tumor cells and frequency of CD107a expression by NKG2A^+^ CD8 T cells that is restored by monalizumab upon co-culture with HLA-E^+^ K562. Paired t-tests were used in (B), (C), (D) and Spearman correlation in (E). Corrections for multiple comparisons were applied in (B) and (C). Only the significant observations (p<0.05) are displayed in (C). * p<0.05, ** p<0.01, *** p<0.001, **** p<0.0001.

We therefore used this experimental model to delineate the effects of NKG2A and PD-1 acquisition on CD8 T cell phenotypes and response to HLA class I-deficient tumors. (Figure 6C, Figure S6D-E). CD49a and CD69 were upregulated upon PD-1 acquisition, while CD103 was specifically upregulated upon further NKG2A acquisition, confirming enhanced TRM features in NKG2A^+^ CD8 T cells. Expression of TFs associated with exhaustion (T-bet, Eomes, TOX) as well as NKG2D and DNAM-1 were progressively increased during TGF-β1-induced CD8 T cell differentiation. TGF-β1-induced NKG2A acquisition on PD-1^+^ CD8 T cells correlated with stronger recent proliferation (KI-67) and production of IFN-γ and XCL1 in response to HLA-deficient K562 tumors (Figure 6C, S6E-F). These results confirm the presence of an alternative NKG2A^+^ exhaustion program on CD8 T cells with enhanced proliferation, tissue-resident memory features and capacity to respond to HLA class I-deficient tumor cells.

### HLA-E-expressing bladder tumors restrict NKG2A^+^ CD8 T cell functions, which are restored upon NKG2A blockade

Previous studies have demonstrated that HLA-E impairs NKG2A^+^ CD8 T cell and NK cell antitumor functions in patients with head and neck squamous cell carcinomas (HNSCC), with promising applications for NKG2A blockade immunotherapy (André *et al.*, 2018). Yet, the effect of HLA-E on TCR/CD28-independent NK-like functions by NKG2A^+^ CD8 T cells remains unknown. We therefore isolated CD8 T cells from bladder tumors (n=6), expanded them with CD3/CD28 tetramer and low-dose IL-2, IL-7, IL-15 for 11-20 days. We then cultured CD8 T cells with K562 tumors that were stably transfected with HLA-E but still deficient for expression of HLA-A, -B, and -C. The HLA-E^+^ K562 cell line was used as a surrogate due to limited numbers and variable HLA-E expression of autologous tumors. We observed that NKG2A blockade (monalizumab) enhanced NKG2A^+^ CD8 T cell degranulation in response to HLA-E^+^ K562 (Figure 6D). NKG2A blockade did not have any effect in the absence of HLA-E (Figure S6G). Additionally, NKG2A blockade effects were restricted to NKG2A^+^ CD8 T cell degranulation, with no effect on their cytokine or chemokine production (Figure S6H).

We demonstrated that HLA-E expression is globally downregulated on primary bladder tumors compared to their immune counterparts, yet there were still residual levels of HLA-E expressed by tumors, which were highly variable across patients (0-96%, mean 17%) (Figure 2C). In this analysis, we observed that the restoration of NKG2A^+^ CD8 T cell degranulation by NKG2A blockade therapy when co-cultured with HLA-E^+^ K562 directly correlates with HLA-E expression on the primary tumors from which these CD8 T cells were isolated (Figure 6E). Collectively, our results demonstrate that HLA-E inhibits NKG2A^+^ CD8 T cell antitumor responses to HLA-ABC-deficient tumors. These data further suggest that NKG2A^+^ CD8 T cell TCR-independent function is impaired by HLA-E. This dysfunction is however partly reversible through use of NKG2A-blockade therapy.

### Increased abundance of CD8 T cells expressing KLRC1 in bladder tumors associates with better response to PD-L1 blockade immunotherapy

Loss of antigen presentation by tumor cells is a common mechanism of tumor evasion in response to PD-1/PD-L1 blockade immunotherapy. NKG2A^+^ PD-1^+^ CD8 T cells being able to degranulate and release cytotoxic mediators in response to HLA-ABC-deficient tumor cells, we evaluated the effect of *KLRC1* (NKG2A) expression in bladder tumors on the response to PD-L1 blockade immunotherapy. We found, in the IMvigor210 cohort of patients with muscle-invasive or metastatic bladder cancer, that *KLRC1*^high^ patients displayed better survival following PD-L1 blockade (Figure 7A). This effect was restricted to patients with higher *CD8A* or *PDCD1* gene expression or higher PD-L1 immunohistochemistry score (Figure 7B-D). These findings demonstrate the importance of the alternative NKG2A/HLA-E immune checkpoint axis in overcoming mechanisms of resistance in PD-L1-treated bladder tumors.

**Figure 7.**
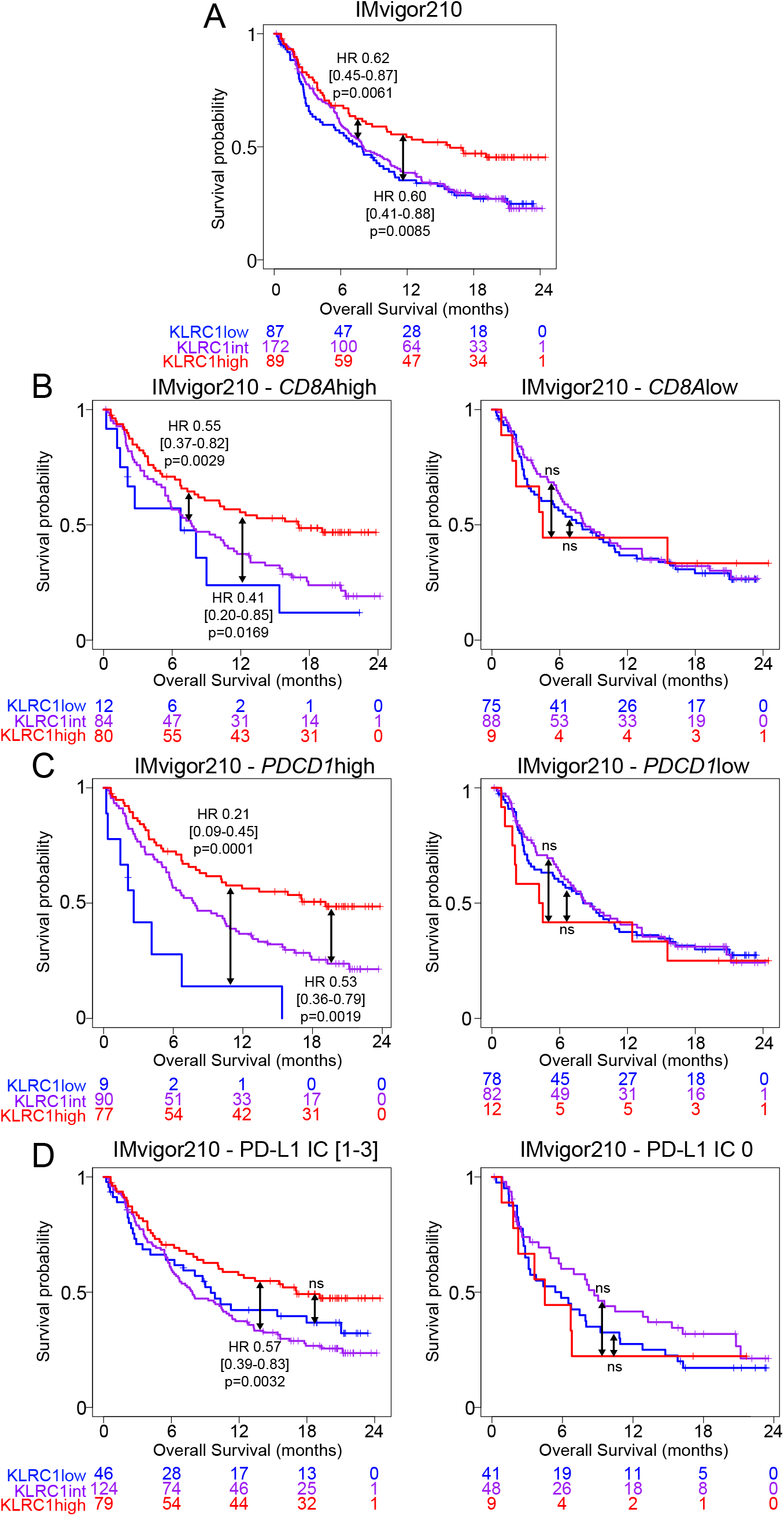
KLRC1 (NKG2A) gene expression correlates with better survival in bladder cancer in response to PD-L1 blockade immunotherapy. Kaplan-meier survival curves of muscle-invasive and metastatic bladder cancer patients treated with PD-L1 blockade immunotherapy from the IMvigor210 cohort, showing the association between *KLRC1* (NKG2A) expression and overall survival in (A) all patients, or patients with (B) *CD8A*^high^ or *CD8A*^low^ (CD8) gene expression, (C) *PDCD1*^high^ or *PDCD1*^low^ (PD-1) gene expression, or (D) a PD-L1 immunohistochemistry score of 0 or [1-3]. Hazard ratios and p-values were calculated using COX proportional hazards regression models. These models were corrected for age, gender and race.

## Discussion

Immunotherapies have revolutionized the treatment of cancer patients, though their efficacy remains variable depending on the tumor indication (Thommen and Schumacher, 2018). The FDA has approved several monoclonal antibodies for blocking PD-1 and PD-L1 in advanced bladder cancer. Yet, this therapeutic strategy yields objective responses in only 15-30% of patients, suggesting additional mechanisms of resistance (Patel et al., 2020). A recent study demonstrated impaired conventional CD8 T cell response in bladder tumors (Han et al., 2021). The HLA-E/NKG2A immune checkpoint axis was recently shown to contribute to CD8 T cell exhaustion. NKG2A blockade strategies have been developed and efficiency confirmed, in particular, for activating CD8 T cells when in combination with anti-PD-L1 antibody blockade therapy in preclinical mouse models (André *et al.*, 2018).

Here, we show that NKG2A is acquired by CD8 T cells alongside other exhaustion markers in bladder tumors. NKG2A is universally defined as an inhibitory receptor. Strikingly, we observed that NKG2A correlates with better survival in MIBC patients with high *CD8A* (CD8) or *PDCD1* (PD-1) expression in their tumor, suggesting a novel alternative immune checkpoint axis that defines enhanced anti-tumor functions by NKG2A^+^ CD8 T cells in bladder tumor settings. We provide the first evidence that NKG2A^+^ CD8 T cells degranulate and secrete cytokines upon binding to NK-activating ligands in the absence of antigen presentation, providing a rationale for their protective role in bladder tumors, where loss of antigen presentation is a common phenotype. We further demonstrated that these TCR-independent, NK-like functions are partly mediated through DNAM-1 interactions with its ligands, CD112 and CD155, expressed on tumors. DNAM-1 expression is significantly higher on NKG2A^+^ CD8 T cells, and DNAM-1-mediated TCR-independent activation is uniquely restricted to the NKG2A^+^ CD8 T cell compartment in healthy individuals. We additionally observed strong downregulation of CD28 on NKG2A^+^ CD8 T cells alongside increased expression of *HSH2D*, the gene encoding AXL which directly inhibits CD28-mediated activation (Shapiro *et al.*, 2004), suggesting a specialization of NKG2A^+^ CD8 T cells toward TCR-independent activation. In support of our findings, a phenotype of CD8 T cell-mediated killing of autologous tumors in the presence of anti-HLA class I antibodies was recently reported (Oh *et al.*, 2020). The study concluded that this observation was reproduced across numerous experiments but without an explanation for the mechanism of action. Our results confirm TCR-independent activation of NKG2A^+^ CD8 T cells in bladder cancer and offer a mechanistic explanation for enhanced anti-tumor functions in response to HLA class I-deficient tumors.

In this study, we also provide the first evidence that the strong downregulation of MHC class I molecules on bladder tumors is significantly pronounced at more advanced, muscle-invasive stages. Known drivers are somatic mutations resulting in the loss of one or more alleles (loss of heterozygosity “LOH”) (Grasso *et al.*, 2018; McGranahan et al., 2017) or post-transcriptional or epigenetic changes in response to the microenvironment (Ritter *et al.*, 2017). We were able to exclude somatic mutations as the dominant driver for the loss in HLA class I expression on bladder tumors based on analyses of the TCGA bladder cohort and through empirical HLA genotyping of matched tumor and germline tissues in our cohort of bladder cancer patients. Thus, it will be important for future studies to investigate epigenetic effects on modulating expression of HLA class I allotypes on tumors.

In cancer (and in chronic viral infection), memory precursor CD8 T cells undergo a transitional exhaustion stage prior to reaching terminal exhaustion. Transitionally exhausted CD8 T cells are defined by stronger effector function, proliferation and response to checkpoint blockade (McLane et al., 2019). This phenotype is lost upon terminal exhaustion. In our analyses, NKG2A^+^ PD-1^+^ CD8 T cells possess stronger evidence of recent proliferation compared to their NKG2A^−^ PD-1^+^ counterparts. *KLRC1*^high^ (NKG2A) CD8 T cells additionally display higher cytolytic content at *mRNA* levels. Further, NKG2A^+^ CD8 T cell TCR-independent function is impaired by HLA-E but restored upon NKG2A blockade. Collectively, these findings suggest that NKG2A^+^ CD8 T cells might represent an additional transitional population, characterized by enhanced proliferation and TCR-independent function that eventually becomes terminally exhausted. Hudson et al. characterized the transcriptome of CD8 T cells upon exhaustion in chronic cytomegalovirus mice models (Hudson et al., 2019). We confirmed in the data from their study that *KLRC1* (NKG2A) (and *MKI67* (KI-67)) expression increases in CD8 T cells upon transitional exhaustion and decreases upon terminal exhaustion, supporting our hypothesis.

We further demonstrated that TRM and exhaustion programs are concomitantly acquired on CD8 T cells in bladder tumors, with CD103 being specifically upregulated upon NKG2A but not PD-1 acquisition. CD103 binds to E-cadherin on tumors, reinforcing the stability of the immunological synapse and enhancing CD8 T cell cytotoxicity (Franciszkiewicz et al., 2013). TRM CD8 T cells have therefore been found to preferentially infiltrate tertiary lymphoid structures within the tumor micro-environment in gastric cancer (Mori et al., 2021). In bladder cancer, TRM CD8 T cells are enriched in bladder tumors compared to the stroma and CD103 expression on CD8 T cells further associates with decreased tumor volume (Wang et al., 2015) and a stronger response to PD-L1 blockade (Banchereau et al., 2021). Thus, stronger TRM features appear to enhance the observed superior anti-tumor functions by NKG2A^+^ CD8 T cells in bladder cancer.

While the TRM program allows NKG2A^+^ CD8 T cells to remain within the tumor tissue and TCR-independent activation contributes to their enhanced anti-tumor functions, HLA-E is a known inhibitor of NKG2A^+^ CD8 T cells and NK cell responses (André *et al.*, 2018). HLA-E is commonly overexpressed on tumor cells, in part due to IFN-γ and senescence pathways (Pereira et al., 2019; van Hall *et al.*, 2019). We demonstrate that HLA-E expression is also strongly downregulated on muscle-invasive bladder tumors, suggesting regulation mechanisms in response to the immunosuppressive environment. Association between tumor progression and MHC class I downregulation was observed for HLA-ABC but not for HLA-E, indicating supplemental sources for HLA-E variation, such as treatment (or expression of the non-classical HLA-F and HLA-G). We show that HLA-E expression on tumors predicts *in vitro* efficacy of NKG2A blockade on the TCR-independent CD8 T cell response. HLA-E^+^ bladder tumors therefore induce NKG2A^+^ CD8 T cell inhibition, that is reversible upon NKG2A blockade. We believe we have defined both a tumor indication in bladder cancer as well as a biomarker of selection by screening for proportion of tumor-infiltrating NKG2A^+^ CD8 T cells and expression of HLA-E on bladder tumors. Future clinical trials may investigate if HLA-E expression on bladder tumors could predict patient response to NKG2A-blockade in combination with PD-(L)1-blockade and, therefore, contribute to personalizing bladder cancer patient treatment.

There are limitations to our study. Mass cytometry is a high-throughput technique that yields reproducible results of strong quality, though requires a large quantity of cells. Therefore, CD8 T cell expansion from bladder tumor specimens, as recently described (Oh *et al.*, 2020), was required in order to apply mass cytometry to our functional assays. We also used the K562 tumor cell line as a surrogate for MHC class I-deficient autologous bladder tumor cells. K562 cells lack endogenous expression of HLA class I and are routinely used to evaluate TCR/MHC-independent activation pathways. Wild-type or HLA-E^+^ K562 cells lack PD-L1 expression and did not afford us the opportunity for studying the combined effects of NKG2A and PD-L1/PD-L1 blockade. An alternative method to overcome these limitations in future studies may be the development of organoids derived from MHC class I-deficient autologous tumor cells.

Finally, we show that *KLRC1* (NKG2A) expression associates with better response to PD-L1 blockade (atezolizumab) in *CD8A*^high^ or *PDCD1*^high^ bladder tumors. Loss of antigen presentation is a common mechanism of resistance to PD-1/PD-L1 blockade (Lee et al., 2020). NKG2A^+^ CD8 T cell TCR-independent functions might therefore contribute, indirectly, to overcoming mechanisms of resistance to anti-PD-1/PD-L1 blockade. While PD-1 blockade specifically restores TCR signaling, NKG2A blockade restores NKG2A^+^ CD8 T cell TCR-independent function. It has indeed been reported that NKG2A prevents the translocation of lipid rafts to the immune synapse and proper amplification of T cell and NK activating signals (Cichocki and Miller, 2019). Our data demonstrate that HLA-E is expressed at low but variable levels on bladder tumors; When expressed, it directly inhibits NKG2A^+^ CD8 T cell response to HLA-ABC-deficient tumors. Given the critical role of NKG2A^+^ CD8 T cells in enhancing the anti-tumor immune response to PD-L1 blockade, combining PD-1/PD-L1 blockade with NKG2A blockade might therefore be worth considering in patients with HLA-E-expressing bladder tumors. Currently, a Phase 1/2 clinical trial of monalizumab in combination with durvalumab is underway in metastatic microsatellite-stable colorectal cancer and will provide preliminary evidence on the safety of this therapeutic strategy (NCT02671435, Communication of Cho et al. at the EMSO 2019 Congress). Collectively, our data indicate that NKG2A^+^ CD8 T cells display a strong capacity for TCR-independent activation that enables them to circumvent tumor escape mechanisms and call for a thorough reappraisal of current protocols that assess CD8 T cell exhaustion and the strategies for restoring their antitumor functions.

## ACKNOWLEDGMENTS

We thank Miriam Merad (Precision Immunology Institute, Icahn School of Medicine at Mount Sinai) for critically reviewing the manuscript; Deepta Bhattacharya ́(University of Arizona) for kindly providing HLA-E^+^ K562 tumors. We acknowledge the expertise and assistance of the Dean’s Flow Cytometry Center of Research Excellence at Mount Sinai. HLA genotyping was funded in whole or in part in Mary Carrington’s lab with federal funds from the Frederick National Laboratory for Cancer Research, under Contract No. HHSN261200800001E, and was supported in part by the Intramural Research Program of the NIH, Frederick National Lab, Center for Cancer Research. The content of this publication does not necessarily reflect the views or policies of the Department of Health and Human Services, nor does mention of trade names, commercial products, or organizations imply endorsement by the U.S. Government. Antibody blocking reagents for NKG2D, NKp30, and NKp46 were provided by the E.M. lab, and the work was supported by Compagnia di San Paolo (2019.866) and Fondazione Associazione Italiana per la Ricerca sul Cancro (AIRC 5 × 1000-21147). The N.B. lab was supported by funding from the Department of Defense Peer Review Cancer Research program Translational team Award No. W81XWH1910269 and from the Parker Institute for Cancer Immunotherapy No. AGR-11611SOW1. The A.H. lab was supported by funding from P30CA196521 and RAI130760A.

## AUTHOR CONTRIBUTIONS

B.S., N.B., and A.H. conceived the project. B.S., N.B., and A.H. conceived the experiments. J.P.S., R.M., P.W., M.G., N.T., and R.B. provided access to the human samples. E.M. provided reagents. B.S., J.D. and A.C. performed the experiments. B.S., A.H., N.B. analyzed the data. J.Z., M.V., T.O.D and D.C performed the LOH HLA and FACETS analyses. A.M.F, J.Z., K.G.B, L.W, R.P.S, S.S, Y-C.W, Y.A.W. collected the RNA sequencing data. D.G., G.K., R.M.d-R, B.L., S.K-S. acquired sample data using the mass cytometer. T.H.T, M.G-B, R.B., E.H-S, E.M.M, R.F-G, M.S performed the imaging experiments. M.C, Y.Y, M.M performed the HLA genotyping analysis. C.H., R.B., S.M. performed analysis of *ImVigor 210* data. A.O.K., E.M., I.M., K-J.M., and P.R. provided intellectual input. B.S., N.B., and A.H. wrote the manuscript.

## STAR METHODS

### RESOURCE AVAILABILITY

#### Lead contact

Further information and requests for resources and reagents should be directed to and will be fulfilled by the Lead Contact Amir Horowitz.

#### Materials availability

This study did not generate new unique reagents.

#### Data and code availability

The single-cell RNA sequencing and mass cytometry data generated in this article will be publicly available.

The algorithms used in this study will be made available at https://github.com.

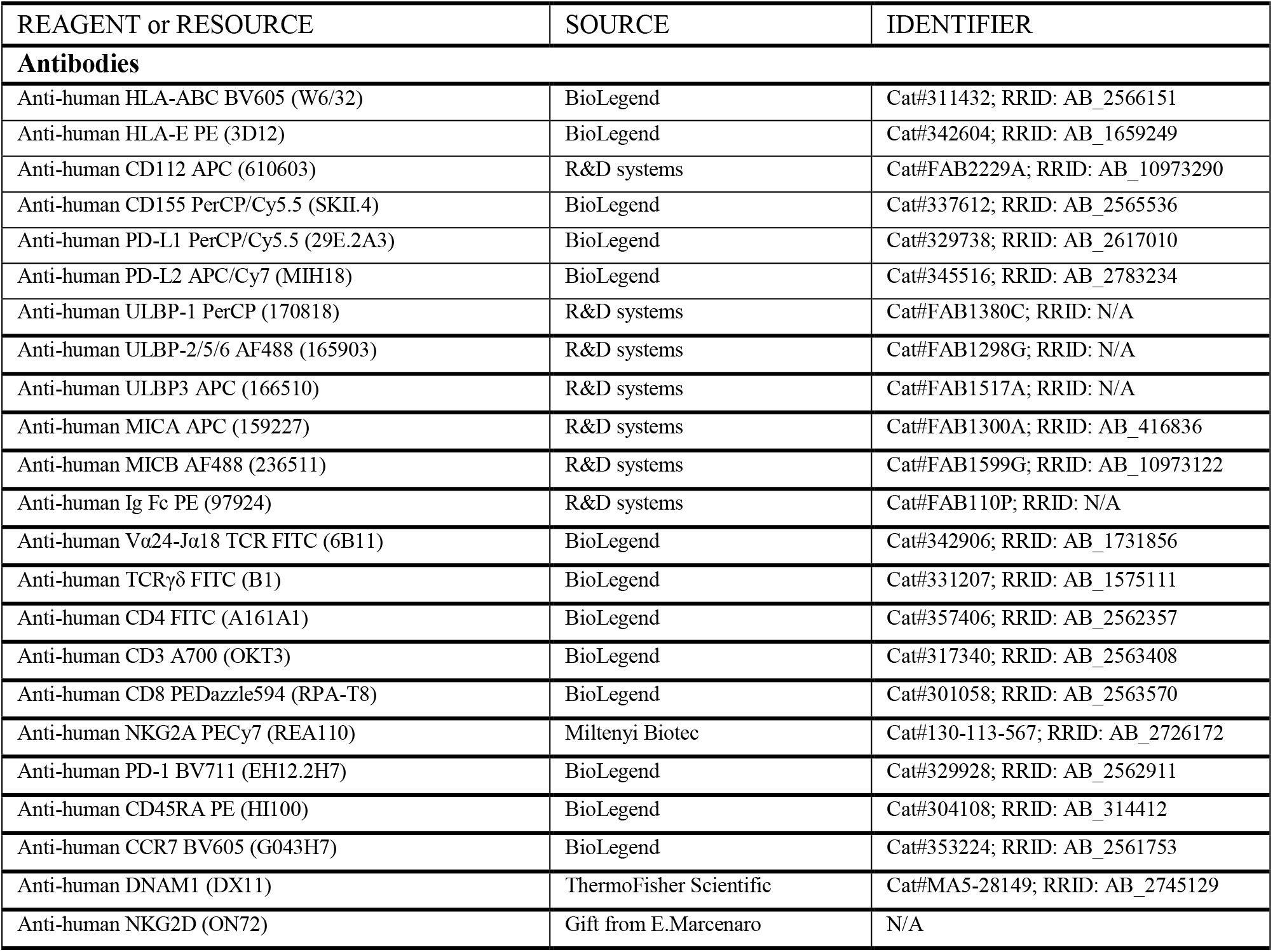
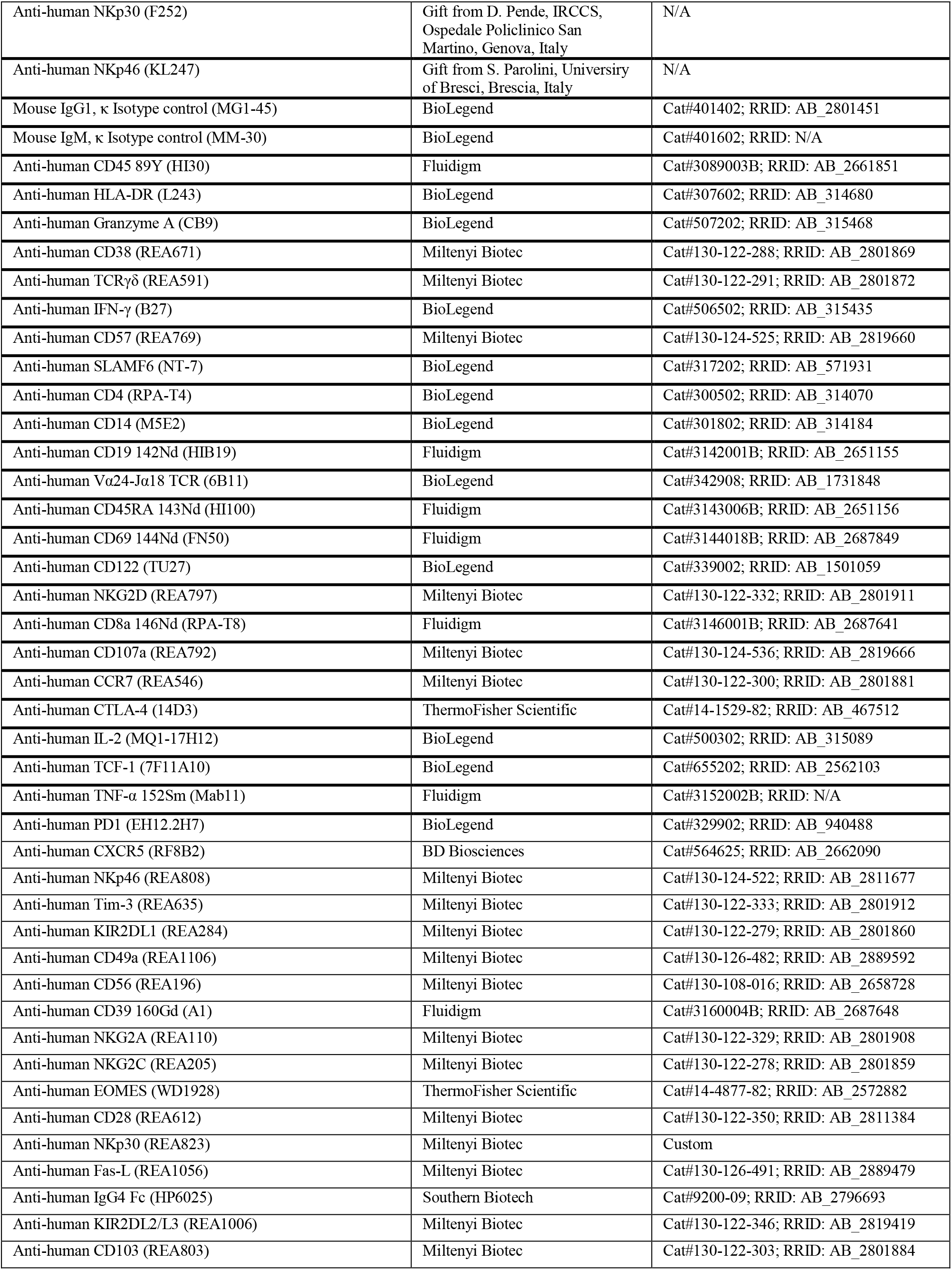
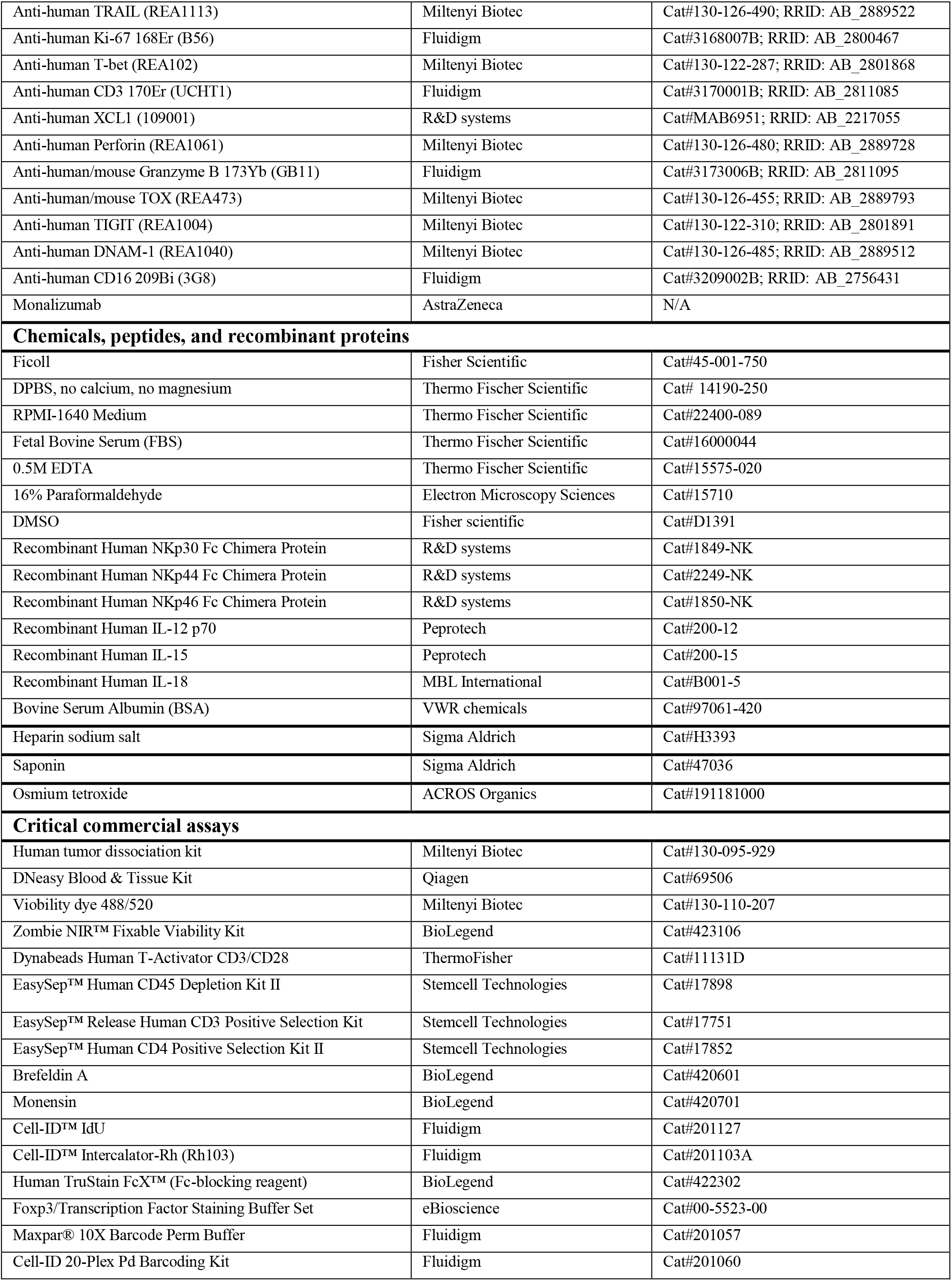
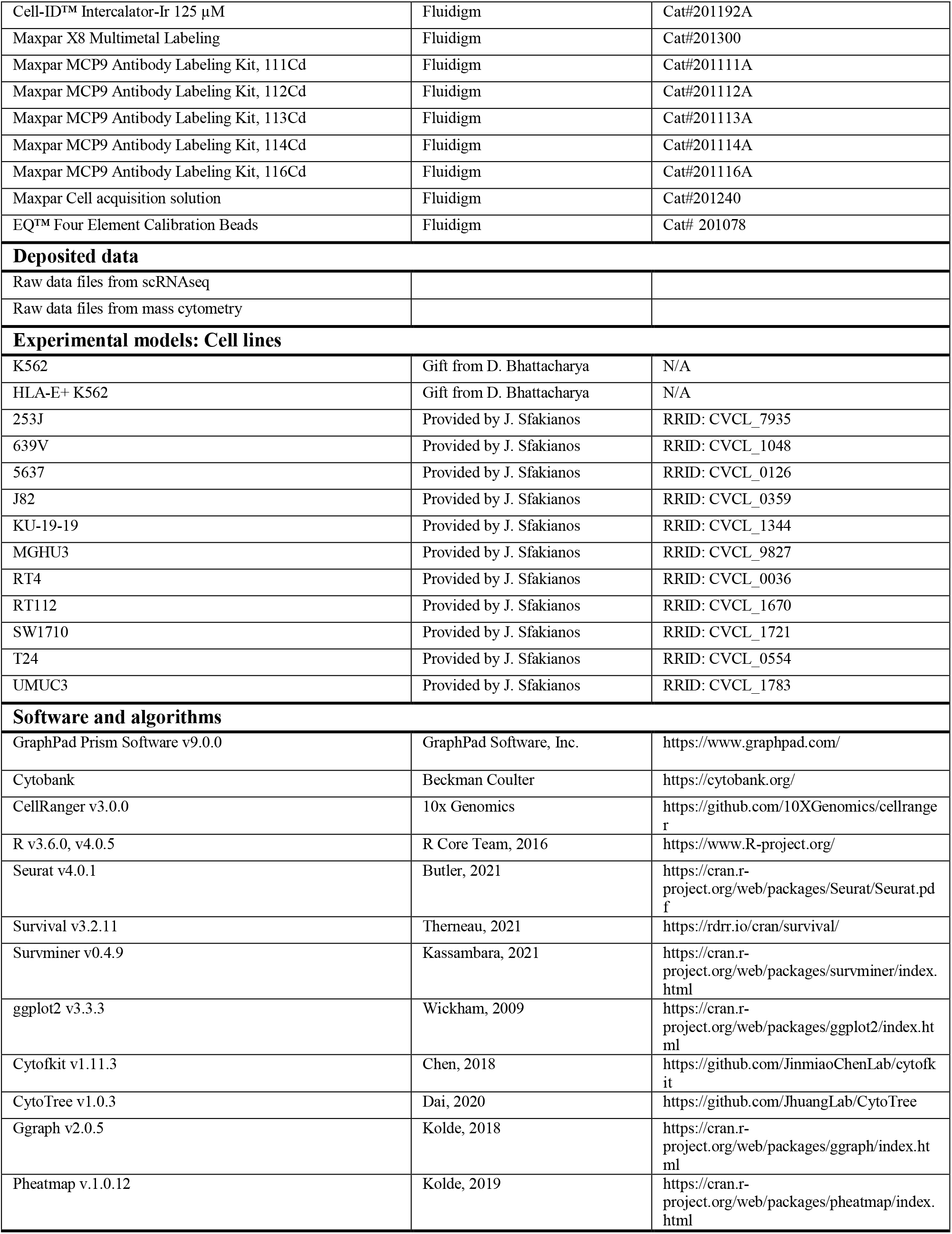
KEY RESOURCE TABLE

### EXPERIMENTAL MODEL AND SUBJECT DETAILS

#### Human subjects

Healthy donor buffy coats were purchased from the New York City Blood Center (New York, NY, USA). Peripheral blood, tumor tissue, adjacent non-tumoral tissue and bladder-draining lymph nodes were obtained from bladder cancer patients upon cystectomy or transurethral resection of bladder tumor (TURBT) at the Mount Sinai Medical Center (New York, NY, USA) after getting informed consent. All protocols were reviewed and approved by the Institutional Review Board at the Icahn School of Medicine at Mount Sinai (IRB 10-1180). Patient information is detailed in Table S1.

#### Cell lines

HLA-E+ and Wild-type K562 cell lines were kindly provided by Deepta Bhattacharya and propagated as recently described (Berrien-Elliott et al., 2020). HLA-E^+^ K562 cells were generated using the AAVS1-EF1a donor plasmid containing the coding sequence for human HLA-E. The K562 were electroporated using a Bio-Rad Gene Pulse electroporation system. HLA-E^+^ cells were sorted to >98% purity.

Bladder cancer cell lines were provided by John Sfakianos: 253J(RRID: CVCL_7935), 639V(RRID: CVCL_1048), 5637(RRID: CVCL_0126), J82(RRID: CVCL_0359), KU-19-19(RRID: CVCL_1344), MGHU3(RRID: CVCL_9827), RT4(RRID: CVCL_0036), RT112(RRID: CVCL_1670), SW1710(RRID: CVCL_1721), T24(RRID: CVCL_0554), UMUC3(RRID: CVCL_1783).

### METHOD DETAILS

#### Sample processing

Ficoll separation was used to obtain PBMCs from healthy donor buffy coats and peripheral blood from bladder cancer patients. Tumor and adjacent, non-involved tissues were digested at 37C using tumor dissociation enzymes (Miltenyi, 130-095-929) and a GentleMACS machine (program 37C_h_TDK_2). Lymph nodes were mechanically dissociated at room temperature using a GentleMACS machine (program m_spleen_04_02). Tumor-, adjacent tissue- and lymph node-derived cell suspensions were then sequentially filtered through 100μM, 70μM and 40μM cell strainers, washed twice with phosphate-buffered saline (PBS) and resuspended in RPMI-1640 medium supplemented with 10% heat-inactivated fetal bovine serum (FBS), 1% Penicillin, 1% Streptomycin and 1% L-glutamine. Cells were counted using a hemocytometer. In certain cases, cells were frozen in FBS supplemented with dimethyl-sulfoxide (DMSO).

#### Flow cytometry

Cells were stained extracellularly for 30 minutes at 4C in FACS buffer (PBS supplemented with 2% heat-inactivated FBS and EDTA 2m*M*). Following the extracellular staining, cells were washed in PBS and incubated during 20 minutes with a viability dye. Cells were then washed with PBS and resuspended in 2% paraformaldehyde prior to their acquisition at an LRS-Fortessa machine (BD Biosciences). Data were analyzed using the Cytobank software. When staining for HLA-E and HLA-ABC, cells were first stained 20 minutes with HLA-E, prior to staining with additional antibodies, including HLA-ABC, due to cross-reactivity between clones 3D12 and W6/32. When staining for NKp30, NKp44 and NKp46 ligands, cells were first stained with NKp30-, NKp44- or NKp46-Fc for 30 minutes on ice, washed with FACS buffer and further stained with anti-human IgG Fc during 30 minutes on ice, prior to incubation with a viability dye.

#### K562 stimulation of CD8 T cells from HD PBMCs

HD PBMCs were thawed and resuspended in cell medium (RPMI-1640 medium supplemented with 10% heat-inactivated FBS, 1% Penicillin, 1% Streptomycin and 1% L-glutamine). CD4^+^ cells were removed using EasySep CD4 positive selection kit (Stemcell, cat#17852) and CD3^+^ cells further enriched using EasySep Release CD3 positive selection kit (Stemcell, cat#17751). Enriched CD8 T cells were then cultured overnight in cell medium supplemented with IL-12, IL-15 and IL-18 (each at 10ng/mL). On the following day, CD8 T cells were co-cultured with K562 at a 1:1 ratio during 5 hours in the presence of brefeldin and monensin. For experiments using blocking antibodies, CD8 T cells were incubated during 30 minutes at room temperature with monalizumab, anti-DNAM1 (DX11), anti-NKG2D (BAT221), anti-NKp30 (F252), anti-NKp46 (KL247), or isotype controls prior to the K562 co-culture.

#### CD3/CD28 stimulation of CD8 T cells from HD PBMCs

HD PBMCs were thawed and resuspended in cell medium (RPMI-1640 medium supplemented with 10% heat-inactivated FBS, 1% Penicillin, 1% Streptomycin and 1% L-glutamine). CD4^+^ cells were removed using EasySep CD4 positive selection kit (Stemcell, cat#17852) and CD3^+^ cells further enriched using EasySep Release CD3 positive selection kit (Stemcell, cat#17751). Enriched CD8 T cells were then cultured for 24h in cell medium supplemented with IL-7 10ng/mL, IL-15 10ng/mL, IL-2 10IU/mL with or without CD3/CD28 Dynabeads and flow cytometry performed at the end of the assay.

#### Expansion of CD8 T cells from bladder cancer patients

CD4^+^ cells were removed from freshly dissociated bladder-draining lymph nodes or bladder tumors using EasySep CD4 positive selection kit (Stemcell, cat#17852) and CD3^+^ cells further enriched using EasySep Release CD3 positive selection kit (Stemcell, cat#17751). Enriched CD8 T cells were seeded at a concentration of 1 million cell/mL in U-bottom 96-well plates in Immunocult medium (Stemcell), in the presence of IL-2 (10IU/mL), IL-7 (10ng/mL), IL-15 (10ng/mL) and CD3/CD28 tetramer (25μL/mL). Half of the medium was replaced with IL-2, IL-7, IL-15 at twice final concentration every 2 days. Cells were split when reaching affluence. CD3/CD28 tetramer was added at Day 10 or Day 11 and eventually at Day 15 expansions longer than 15 days. CD8 T cells from bladder-draining lymph nodes were expanded for 8-13 days (8, 11, 12, 13, 13, 13 days), while CD8 T cells from bladder tumors were expanded for 11-17 days (11, 11, 11, 14, 15, 17 days).

CD8 T cells from bladder tumors were directly co-cultured upon expansion for 5 hours with monensin, brefeldin +/− WT (HLA-E^−^) or HLA-E^+^ K562 (1:1 E:T ratio). When using NKG2A blockade, CD8 T cells were stained with monalizumab for 30 minutes at room temperature prior to co-culture.

CD8 T cells from bladder-draining lymph nodes were FACS-sorted based on CD49a, NKG2A and PD-1 upon 8-13-day expansion with a purity higher than 95% (BD FACSAria II, 100μM nozzle). Resulting cell populations were cultured in Immunocult medium (Stemcell) in the presence of IL-2 (10IU/mL), IL-7 (10ng/mL), IL-15 (10ng/mL), CD3/CD28 tetramer (25μL/mL) +/− TGF-β (5ng/mL) for another 3 days, with replacement of half medium on the second day. Mass cytometry stainings were performed at day two and day three. On day three, cells were washed, counted, and cultured for 5h in the presence of monensin, brefeldin +/− WT (HLA-E^−^) K562 (1:1 E:T ratio) prior to staining.

#### Mass Cytometry antibody preparation

When available, conjugated antibodies were purchased from Fluidigm. Remaining antibodies were purchased carrier-free and were conjugated in-house using the Maxpar X8 and MCP9 labeling kits (Fluidigm), as per the manufacturer instructions. Platinum barcodes were prepared as described previously (Hartmann et al., 2018). All antibodies were titrated prior to use.

#### Mass Cytometry staining

Cells were incubated during 20 minutes at 37C in cell medium (RPMI-1640 medium supplemented with 10% heat-inactivated FBS, 1% Penicillin, 1% Streptomycin and 1% L-glutamine) in the presence of IdU (Fluidigm, cat#201127), Rh103 (Fluidigm, cat#201103A) and eventually anti-IgG4 if cells were co-cultured with K562 and monalizumab. Cells were centrifuged and washed with PBS 0.2% Bovine Serum Albumin (BSA) prior to a 3-minute incubation on ice with an Fc-blocking reagent (BioLegend, cat# 422302). Samples were then washed and incubated with Platinum barcodes for 30 minutes on ice. When staining a maximum of 4 samples, cells were single-barcoded using 194Pt, 195Pt, 196Pt or 198Pt. When staining 5 or 6 samples, cells were stained with a dual combination of these barcodes. Cells were then washed with PBS 0.2% BSA, barcoded samples pooled together, washed and stained with extracellular antibodies for 30 minutes on ice in PBS 0.2% BSA. Samples were then washed with PBS 0.2% BSA and resuspended in Fixation/Perm buffer (Invitrogen, cat#00-5523-00) for 30 minutes on ice. Cells were then centrifuged and washed with Maxpar Barcode Perm Buffer (Fluidigm, cat#201057) and barcoded using the Cell-ID 20-Plex Pd Barcoding kit (Fluidigm, cat#201060). Barcoded samples were then washed with permeabilization buffer (Invitrogen, cat#00-5523-00) and pooled together prior to intracellular staining in permeabilization buffer in the presence of Heparin 100U/mL for 30min on ice. Stained cells were washed with permeabilization buffer and resuspended in PBS in the presence of PFA 2.4%, saponin 0.08%, Osmium tetroxide 0.075nM and Ir 0.125uM (Fluidigm, cat#201192A). Stained samples were finally washed and resuspended in PBS 0.2% BSA and data acquired within four days, or frozen in FBS/DMSO 90/10. Antibody panels 1-3 were used to stain HD PBMCs (panel 1: Figure3A-B, panel 2: Figure3C, panel 3: Figure3D-G). Antibody panel 3 was used to stain bladder cancer patient samples.

#### Mass Cytometry sample acquisition and processing

Immediately prior to acquisition, samples were washed with Cell Staining Buffer and Cell Acquisition Solution (Fluidigm) and resuspended in Cell Acquisition Solution at a concentration of 1 million cells per ml containing a 1:20 dilution of EQ normalization beads. The samples were acquired on the Fluidigm Helios mass cytometer using the wide bore injector configuration at an acquisition speed of < 400 cells per second. The resulting FCS files were normalized and concatenated using Fluidigm’s CyTOF software. The FCS files were further cleaned using the Human Immune Monitoring Center at Mt. Sinai’s internal processing pipeline. The pipeline removes aberrant acquisition time-windows of 3 seconds where the cell sampling event rate goes above or below 2 standard deviations from the mean acquisition speed. EQ normalization beads are removed as well as low DNA intensity events. Samples were demultiplexed by calculating the cosine similarity of every cell’s Palladium barcoding channel to every possible barcode used in a batch. Once the cell has been assigned to a sample barcode, a signal-to-noise metric is calculated by taking the difference between its highest and second highest similarity scores. Any cells with low signal-to-noise are flagged as multiplets and removed from that sample. Finally, acquisition multiplets are removed based on the Gaussian parameters Residual and Offset acquired by the Helios mass cytometer. CD8 T cells were then selected on Cytobank as live CD45^+^ CD4^−^ CD14^−^ CD19^−^ Vα24/Jα18 TCR^−^ γδTCR^−^ CD3^+^ CD8^+^ cells.

#### Mass Cytometry data analysis

Cytobank was used to assess the frequencies of cells expressing markers of interest. For further analyses, CD8 T cells were exported from Cytobank as FCS files and imported into the R software using the cytof_exprsMerge function (Cytofkit). Inverse hyperbolic sine transformation with a cofactor of 5 was applied. For bladder cancer patient samples, all cells were analyzed. For healthy donor PBMCs, a maximum of 1,000 NKG2A^+^ and 1,000 NKG2A^−^ CD8 T cells was imported per donor and condition. Associations between phenotype and K562-induced functional response on HD CD8 T cells were determined using linear models (lm() function). Models were constructed as X~Y*Z, with X the expression of a functional marker, Y the expression of a phenotypic marker, and Z the condition (medium or K562). Results were visualized with the ggraph package.

FlowSOM cluster analysis was performed on K562-stimulated HD CD8 T cells using the cytof_cluster function (Cytofkit) based on the percentage of expression of each marker, with an assigned number of clusters (n=15). The threshold of positivity for each marker was defined using manual gating. One anecdotical cluster was removed from the analysis, as containing less than 1% of the total cells. Heatmap of the cluster phenotypes was obtained using the pheatmap function. Functional strength indexes were calculated for each cluster and condition as the sum of all functional marker strength indexes. Functional marker strength indexes were defined as [frequency of cells expressing a marker X]*[mean signal intensity of the marker X on X^+^ cells].

T-SNE analysis on *ex vivo* bladder samples was conducted using the cytof_dimReduction function (Cytofkit) on all CD8 T cells from the tumor and adjacent tissue (n=1,089 and 564, respectively) and on a maximum of 1,100 CD8 T cells from the bladder-draining lymph nodes (maximum number of iterations = 1,000). Results were visualized using the ggplot2 and pheatmap packages.

Pseudotime was calculated on *ex vivo* bladder tumors using the CytoTree package. Cells were clustered, principal component analysis was performed, cell dimensionality was reduced based on UMAP and t-SNE methods and trajectory trees built. Root clusters were manually defined and pseudotime was calculated using the t-SNE dimensions. Results were visualized with the plot2D, plotTree, plotPieTree and plotPseudotime functions. Spearman correlations were performed on marker expression levels and pseudotime. P-values were adjusted using the Benjamini-Hochberg method and results represented with the ggplot2 package.

Pseudotime was determined using the same pipeline on expanded CD8 T cells from bladder tumors that were cultured for 5 hours with or without K562. Pseudotime effect on CD8 T cell function was assessed on TGF-β-cultured CD8 T cells using linear models (lm() function). Models were constructed as X~Y*Z, with X the expression of a functional marker, Y the pseudotime and Z the condition (medium or K562). Effect of the expression of each marker on CD8 T cell function was assessed using linear models on TGF-β-cultured CD8 T cells as X~Y*Z, with X the expression of a functional marker, Y the expression of a phenotypic marker and Z the condition (medium or K562). P-values were adjusted using the Benjamini-Hochberg method. Results were visualized using the GraphPad Prism software.

#### TCGA and IMvigor analyses

The TCGA cohort contains 406 total patients, including 371 MIBC and n=5 NMIBC patients. Pathologic T stage information could not be retrieved for the 30 remaining patients. Nine patients were previously treated with radiation, one with intravesical BCG and radiation, and 36 with intravesical BCG alone. Treatment information was not accessible for remaining patients. *KLRC1*, *CD8A* and *PDCD1* gene expression levels were scaled prior to all analyses. The 25% and 75% quantiles were used to distinguish *KLRC1*^low^ (<25%), *KLRC1*^int^ (25-75%) and *KLRC1*^high^ (>75%) patients. The median was used to identify *CD8A*^low^/*CD8A*^high^ and *PDCD1*^low^/*PCDD1*^high^ patients. Patients from the IMvigor cohorts were additionally divided based on the PD-L1 score assessed by immunohistochemistry. Analyses were performed using the survminer and survival packages. Survival curves were generated with Kaplan-Meier analyses. Hazard ratios and p-values were calculated with Cox proportional hazards regression models corrected for age and number of somatic mutations. Gender has no effect on survival in our model and was therefore not included as a correction factor.

#### Single-cell RNA sequencing sample preparation

ScRNA-seq was performed on freshly dissociated bulk tumor cells (n=7) or on CD45^+^ and CD45^−^ FACS-sorted freshly dissociated tumor cells (n=7)(Table 1) using a Chromium controller (10x Genomics), as previously described(Wang *et al.*, 2021). Briefly, gel-bead in emulsion were generated, cells were lysed and barcoded cDNA amplified for 12 cycles. Amplified cDNA was fragmented and subjected to end-repair, poly A-tailing, adapter ligation, and 10x–specific sample indexing following the manufacturer’s protocol. Bioanalyzer (Agilent Technologies) and QuBit (ThermoFisher Scientific) were used to quantified the libraries, which were then sequenced in dedicated flowcells in paired-end mode on a HiSeq 2500 (Illumina) targeting a depth of 5E10^4^–1E10^5^reads per cell.

#### Single-cell RNA sequencing analysis

Raw sequencing data were aligned and quantified using CellRanger against the provided GRCh38 human reference genome. Seurat was then used for all remaining steps. For each sample, cells were first selected as expressing less than 16-20% mitochondrial genes and displaying a minimum of 200-300 and a maximum of 2500-3500 features. Data were then log-normalized using a scale factor of 10,000. The 2,000 most variable features were then identified, data were scaled based on all the features, and principal component analysis was performed. Dimensionality of the dataset was then assessed using the JackStraw and ElbowPlot functions. Clusters were calculated and data dimensions were reduced using the t-SNE and UMAP methods. Metadata containing clinical information was added to each sample prior to merging. Bulk cells were merged for the HLA analysis, while CD8 T cells only were merged for the CD8 T cell analysis. CD8 T cells were identified using the FindAllMarkers and the VlnPlot functions. CD8 T cell clusters were extracted and any contaminating CD4^+^, CD3D^−^ CD3E^−^ CD3G^−^ or CD8A^−^ CD8B^−^ cell was removed using the WhichCells function. Merged data were then log-normalized using a scale factor of 10,000. As described above, highly variable features were identified, the data were scaled, principal component analysis was performed, and dimensionality of the dataset was determined (n=88 for bulk cell analysis, n=30 for CD8 T cells analysis). The data were then integrated to remove batch effect based on the dataset dimensionality and the most highly variable features (top 3,000 features with the addition of HLA-A, HLA-B, HLA-C and HLA-E genes for bulk cells analysis and top 10,000 genes for CD8 T cell analysis). The integrated data were then scaled, principal component analysis was performed and clusters were identified using the FindClusters function with a 0.5 resolution parameter. T-SNE and UMAP analyses were then performed on the integrated datasets. For HLA analyses, differentially expressed genes were selected using the FindAllMarkers function. Genes expressed by at least 25% of the cells and presenting a log(fold change) of at least 0.25 were selected. Clusters were manually annotated based on these differentially expressed genes. The FeaturePlot function was used to display HLA-A, -B, -C, -E and B2M expression on the UMAP clusters. Differential expression of HLA-A, -B, -C, -E and B2M between the tumor and immune clusters was assessed for each donor on the RNA assay using the FindMarkers function. For CD8 T cell analyses, clusters were manually annotated based on differentially expressed genes that were selected on the integrated assay as expressed by at least 25% of the cells and displaying a log(fold change) of at least 0.05. KLRC1^high^ vs KLRC1^low^ cells were identified within each cluster based on the median integrated gene expression level. Heatmaps were obtained with the DoHeatmap function (Seurat), violin plots with the VlnPlot function (Seurat) and dot plots with the ggplot2 package.

#### DNA extraction and HLA genotyping

DNA was extracted from bulk PBMCs, bulk tumors or the CD45^−^ tumor fraction using the DNeasy blood and tissue kit (Qiagen). The CD45^−^ tumor fraction was obtained using the EasySep human CD45 depletion kit (Stemcell technologies). DNA was quantified using a Nanodrop spectrophotometer (Thermofisher) and stored at −20C. *HLA class I* genotyping was performed by M.Carrington’s laboratory at the National Cancer Institute (Bethesda, MD, USA), following a previously described protocol (Bashirova et al., 2020).

#### TCR clonality analysis

TCR clonality in *KLRC1*^high/low^ CD8 T cells from bladder tumors (n=7) and non-involved adjacent tissue (=2) was determined using previously published data (Oh *et al.*, 2020). ScRNAseq data was downloaded and pre-processed following our scRNAseq analysis pipeline. *KLRC1* threshold for high versus low expression was manually assigned as 1 based on scaled gene expression. *KLRC1* expression (high vs low) per cell was then combined with alpha and beta CDR3 information to determine the frequency of TCRs expressed by 1, 2 or at least 3 cells in the *KLRC1*^high^ or *KLRC1*^low^ fractions.

### QUANTIFICATION AND STATISTICAL ANALYSIS

Details on the statistic tests used can be found in the figure legends. Statistical analyses were performed using R software version 3.6.0 or 4.0.5 and GraphPad Prism software version 9.0.0. Bar graphs show mean ± standard error mean. Statistical significance is defined as p-value <0.05. P-values are indicated in the figure legends and on the plots. * p<0.05, ** p<0.01, *** p<0.001, **** p<0.0001.

