## Supplemental data for "NKG2A and HLA-E define a novel alternative immune checkpoint axis in bladder cancer"

#### SUPPLEMENTAL TABLE AND FIGURE TITLES AND LEGENDS

**Table S1 Single-cell RNA sequencing cluster assignment, Related to Figure 2**
(Provided as an Excel file)

**Table S2 HLA class I genotype of bladder cancer patients, Related to Figure 2**

**Figure S1. Bladder tumor cell-surface expression of HLA class I and ligands for PD-1 and activating NK cell receptors, Related to Figure 2**
The expression of HLA class I and ligands for PD-1 (PD-L1/-L2), DNAM-1 (CD112, CD155), NKG2D (ULBPs, MICA, MICB), NKp30 ("NKp30-L"), NKp44 ("NKp44-L") and NKp46 ("NKp46-L") was assessed by flow cytometry on n=11 bladder cancer lines from histological grade 1 to grade 3 as well as on the control cell line K562. Grey: unstained control, Red: stained cells.

**Figure S2. NKG2A-expressing CD8 T cells are differentiated and possess TCR-independent NK-like functions in healthy individuals, Related to Figure 3**
(A-D) Mass cytometry was performed *ex vivo* on HD PBMCs (n=20).
(A) CMV and gender effect on NKG2A expression on CD8 T cells from HD PBMCs depending on differentiation stage.
(B) Correlation between age and NKG2A expression on CD8 T cells depending on differentiation stage.
(C) Correlation between HLA-E expression on lymphocytes and NKG2A expression on CD8 T cells depending on their differentiation stage.
(D) NKp30, NKp46, NKG2D and DNAM-1 expression on NKG2A$^{+/-}$ CD8 T cells depending on their differentiation stage.
(E) Representative NKG2A and CD28 staining on CD8 T cells.
(F-H) CD8 T cells were isolated from HD PBMCs and recovered overnight with IL-12, IL-15, IL-18 10ng/mL prior to a 5h co-culture with K562 in the presence of brefeldin and monensin (n=20)
(F) Degranulation, production of IFN-γ, TNF-α, XCL1, IL-2 and cytolytic content of CD8 T cells upon K562 stimulation.
(G) Amount of functional markers acquired or released upon K562 stimulation in CD8 T cells and NK cells.
(H) Effect of CD8 T cell NKG2A expression, lymphocyte HLA-E expression, age, gender and combined effect of NKG2A with HLA-E on the upregulation of functional markers upon K562 stimulation.
(I-J) CD8 T cells were isolated from HD PBMCs and recovered overnight with IL-12, IL-15, IL-18 10ng/mL prior to a 5h co-culture with K562 in the presence of brefeldin and monensin in the presence or absence of anti-DNAM-1 blocking antibody (n=10).
(I) Correlation between NKG2A or CD28 expression within each cluster and Log2(Fold change) of the expression of XCL1 upon K562 stimulation.
(J) Log2(Fold change) of IFN-γ and TNF-α expression upon anti-DNAM-1 blocking antibody addition, versus isotype control, in all clusters.
Pearson correlation was used in (B), (C), (I), paired t-tests in (F), linear models in (H) and unpaired t-tests in (I), (J). All statistical tests were adjusted for multiple comparison. Only the significant comparisons are displayed in (F). "ns" $p>0.05$, * $p<0.05$, ** $p<0.01$, *** $p<0.001$, **** $p<0.0001$.

**Table S3 Mass cytometry panels**

**Figure S3. NKG2A defines an alternatively exhausted subset of TRM CD8+ T cells that retain TCR-independent functions in bladder tumors, Related to Figure 4**
(A-D) Mass cytometry was performed on *ex vivo* bladder tumor-draining lymph nodes, tumor and adjacent tissue from bladder cancer patients.

(A) Summary *ex vivo* expression of cytotoxic mediators on CD8 T cells from bladder tumor-draining lymph nodes (n=5), tumor (n=7) and adjacent tissue (n=6). Lines show matching samples from a unique donor.
(B) Association between normalized mean signal intensity of key markers on CD8 T cells with pseudotime (n=7).
(C) CD8 T cell phenotype in bladder tumor-draining lymph nodes that are tumor-infiltrated ("TILN", n=2) or not tumor-infiltrated ("non-TILN", n=3).
(D) Phenotype of $PD1^{+}$ $NKG2A^{-}$ and $PD1^{+}$ $NKG2A^{+}$ CD8 T cells in the tumors (n=6). Only the non-significant markers are displayed.
Paired t-tests were used in (A), (D).

**Figure S4. *KLRC1* (NKG2A) expression in CD8 T cells does not associate with a skewed TCR clonal repertoire, Related to Figure 4**

(A-D) *KLRC1*$^{low/high}$ CD8 T cell clonality was determined by analyzing publicly available scRNAseq and TCRseq data from bladder tumors or non-involved adjacent tissue(Oh *et al.*, 2020).
(A) Frequency of TCRs present in 1, 2 or at least 3 cells per sample, depending on *KLRC1* expression.
(B) Summary data in tumor and non-involved adjacent tissue.
(B) Summary data in tumor tissue.
(C) Frequency of TCRs present in 1 cell depending on *KLRC1* expression.
(D) Number of TCRs present in at least 2 cells that are unique or common to *KLRC1*$^{low}$ and *KLRC1*$^{high}$ cells per sample. Only samples with at least 2 TCRs present in at least 2 *KLRC1*$^{high}$ cells were selected.

**Figure S5. Compositional makeup of CD8 T cell clusters in bladder tumors according to pathology stage, Related to Figure 5**

(A-B) Single-cell RNA sequencing was performed on *ex vivo* bladder tumors (n=14). Gene expression was integrated to remove batch-effect.
(A) Cluster distribution per donor and tumor stage (n=14). One column represents one patient.
(B) Violin plots of gene expression per cluster.

**Figure S6. TGF-β1 induces NKG2A expression on CD8 T cells and enhances their TCR-independent NK-like anti-tumor activity in response to NKG2A-blockade in bladder cancer, Related to Figure 6**

(A-F) CD8 T cells were isolated from bladder tumor-draining lymph nodes and expanded for 8-13 days with IL-2, IL-7, IL-15 and CD3/CD28 tetramer. $CD49a^{-}$ $NKG2A^{-}$ PD1- (n=5), $CD49a^{+}$ $NKG2A^{-}$ $PD1^{-}$ (n=6), $CD49a^{+/-}$ $NKG2A^{-}$ $PD1^{+}$ (n=6) and $CD49a^{+/-}$ $NKG2A^{+}$ $PD1^{-}$ (n=5) cells were then FACS-sorted and expanded during 3 days with IL-2, IL-7, IL-15 and CD3/CD28 tetramer with or without TGF-β prior to co-culture with K562. Mass cytometry was performed at all timepoints.
(A) Representative staining of NKG2A/PD1 expression on FACS-sorted cells upon 3-day expansion (n=1 patient).
(B) Summary data of NKG2A/PD1 expression on FACS-sorted cells upon 3-day expansion.
(C) Upregulation of functional markers upon K562 co-culture of CD8 T cells that have been FACS-sorted as $CD49a^{-}$ or $CD49a^{+}$ $NKG2A^{-}$ $PD1^{-}$ CD8 T cells and expanded during 3 days with TGF-β.
(D) Expression of markers associated with tissue-residency, exhaustion and NK/CD8T cell function on $CD49a^{+/-}$ $NKG2A^{-}$ $PD1^{-}$ CD8 T cells before and after 3-day expansion with or without TGF-β.
(E) Expression of markers associated with tissue-residency, exhaustion and NK/CD8T cell function alongside NKG2A and PD-1 acquisition upon 3-day expansion of $NKG2A^{-}$ $PD1^{-}$ CD8 T cells with TGF-β.
(F) Expression of functional markers alongside NKG2A and PD-1 acquisition upon 3-day expansion of $NKG2A^{-}$ $PD1^{-}$ CD8 T cells with TGF-β and 5h co-culture with K562.
(G-H) CD8 T cells were isolated from bladder tumors (n=6) and expanded for 11-17 days with IL-2, IL-7, IL-15 and CD3/CD28 tetramer, prior to 5h co-culture with Wild-type (WT) or HLA-$E^{+}$ K562.

(G) Degranulation of NKG2A$^+$ CD8 T cells upon co-culture with WT K562 in the presence or absence of monalizumab.
(H) Production of IFN-γ, TNF-α, XCL1 and IL-2 by NKG2A$^+$ CD8 T cells upon co-culture with HLA-E$^+$ K562 in the presence or absence of monalizumab.
Paired t-tests were used in (B)-(H). P-values were corrected for multiple comparisons in (B), (D), (E), (F). "ns" $p>0.05$, * $p<0.05$, ** $p<0.01$, *** $p<0.001$, **** $p<0.0001$.

**Table S4 CD8 T cell exhaustion genes, Related to Figure 5**
(Provided as an Excel file)

**Table S5 Differentially expressed genes between *KLRC1*$^{high}$ and *KLRC1*$^{low}$ CD8 T cells, Related to Figure 5**
(Provided as an Excel file)

Table S2: HLA class I genotype of bladder cancer patients. Related to figure 2.

| ID | HLA-A_1 | HLA-A_2 | HLA-B_1 | HLA-B_2 | HLA-C_1 | HLA-C_2 |
|---|---|---|---|---|---|---|
| #9 | 24:02 | 25:01 | 41:01 | 51:01 | 15:02 | 17:01 |
| #10 | 02:01 | 02:01 | 38:01 | 48:01 | 08:03 | 12:03 |
| #11 | 02:01 | 31:01 | 35:02 | 52:01 | 04:01 | 12:02 |
| #12 | 23:01 | 26:01 | 35:01 | 53:01 | 04:01 | 06:02 |
| #13 | 11:01 | 24:20 | 35:03 | 48:01 | 04:01 | 08:03 |
| #14 | 01:01 | 23:01 | 53:01 | 81:01 | 04:01 | 18:01 |
| #15 | 02:01 | 32:01 | 15:71 | 18:01 | 03:03 | 07:01 |
| #16 | 02:01 | 02:05 | 35:01 | 58:01 | 04:01 | 07:01 |
| #17 | 01:01 | 26:01 | 35:02 | 38:01 | 06:02 | 12:03 |
| #18 | 11:01 | 24:02 | 13:02 | 52:01 | 06:02 | 12:02 |
| #19 | 02:01 | 02:11 | 35:01 | 49:01 | 04:01 | 07:01 |
| #20 | 02:01 | 23:01 | 42:01 | 44:03 | 04:01 | 17:01 |
| #21 | 23:01 | 26:01 | 35:01 | 53:01 | 04:01 | 06:02 |
| #22 | 01:01 | 02:05 | 35:02 | 41:01 | 04:01 | 07:01 |
| #23 | 02:10 | 11:02 | 40:06 | 55:02 | 01:02 | 08:01 |
| #24 | 02:05 | 23:01 | 44:03 | 49:01 | 04:01 | 07:01 |

Figure S1. Bladder tumor cell-surface expression of HLA class I and ligands for PD-1 and activating NK cell receptors. Related to Figure 2

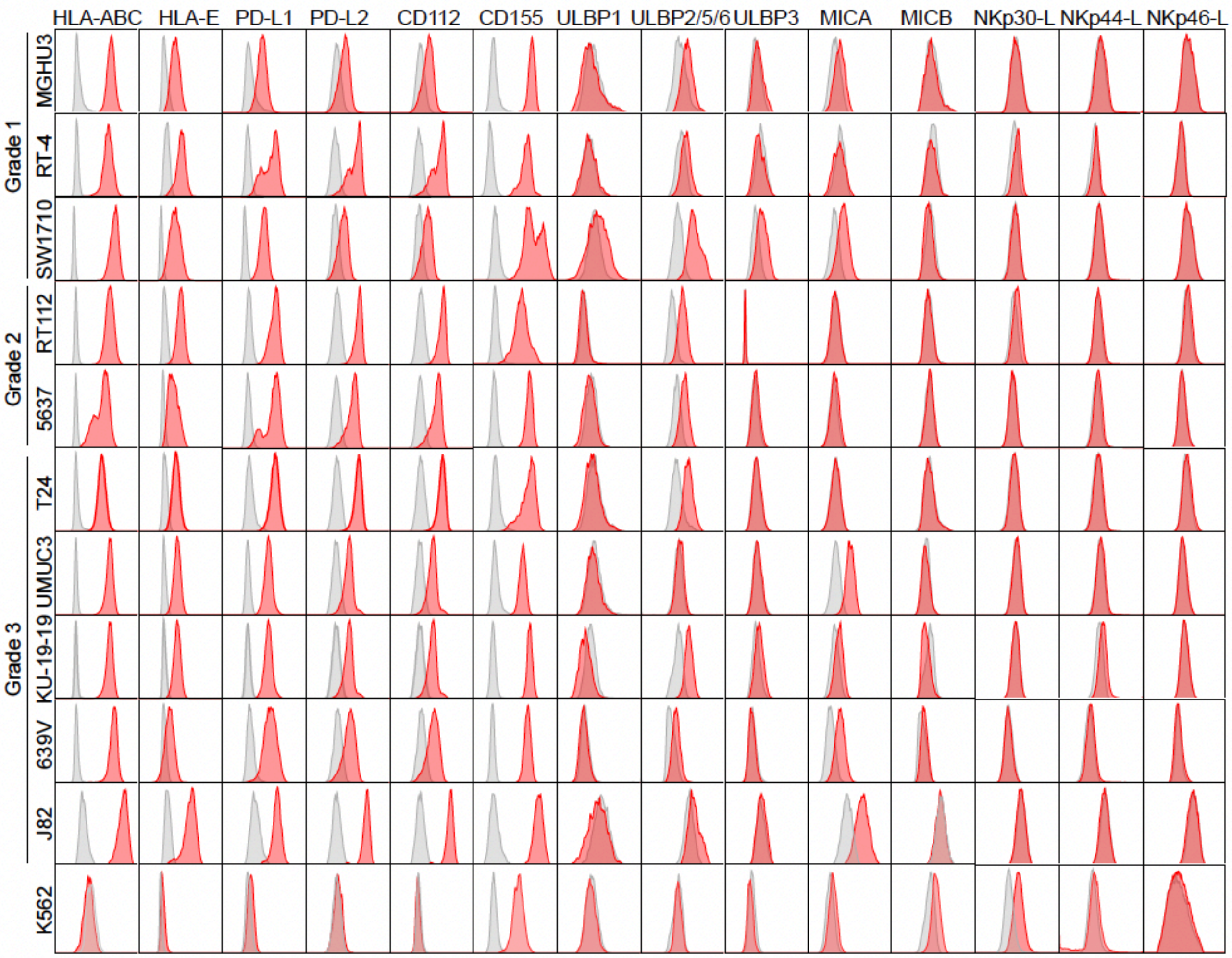

### Table S3: Mass cytometry panels

<table>
<tr><th></th><th>Panel 1</th><th>Panel 2</th><th>Panel 3</th><th>Panel 4</th></tr>
<tr><td>89 Y</td><td colspan="4">CD45</td></tr>
<tr><td>111Cd</td><td colspan="4">HLA-DR</td></tr>
<tr><td>112Cd</td><td colspan="4">Granzyme A</td></tr>
<tr><td>113Cd</td><td colspan="4">CD38</td></tr>
<tr><td>114Cd</td><td colspan="4">γδTCR</td></tr>
<tr><td>115 In</td><td>KLRG1, CRTH2</td><td colspan="3">IFN-γ</td></tr>
<tr><td>116Cd</td><td colspan="4">CD57</td></tr>
<tr><td>141 Pr</td><td></td><td colspan="2">Ksp37</td><td>SLAMF6</td></tr>
<tr><td>142Nd</td><td colspan="4">CD4, CD14, CD19, Vα24 TCR</td></tr>
<tr><td>143 Nd</td><td colspan="4">CD45RA</td></tr>
<tr><td>144 Nd</td><td>CD122</td><td>CD69</td><td>CD122</td><td>CD69</td></tr>
<tr><td>145 Nd</td><td colspan="4">NKG2D</td></tr>
<tr><td>146 Nd</td><td colspan="4">CD8</td></tr>
<tr><td>147 Sm</td><td>NKp80</td><td colspan="3">CD107a</td></tr>
<tr><td>148 Nd</td><td colspan="4">CCR7</td></tr>
<tr><td>149 Sm</td><td colspan="4">CTLA-4</td></tr>
<tr><td>150 Nd</td><td>ROR-γt</td><td colspan="3">IL-2</td></tr>
<tr><td>151 Eu</td><td colspan="4">TCF-1</td></tr>
<tr><td>152 Sm</td><td>CD25</td><td colspan="3">TNF-α</td></tr>
<tr><td>153 Eu</td><td colspan="4">PD1</td></tr>
<tr><td>154 Sm</td><td colspan="4">CXCR5</td></tr>
<tr><td>155 Gd</td><td colspan="4">NKp46</td></tr>
<tr><td>156 Gd</td><td colspan="4">Tim-3</td></tr>
<tr><td>158 Gd</td><td colspan="3">KIR2DL1</td><td>CD49a</td></tr>
<tr><td>159 Tb</td><td colspan="4">CD56</td></tr>
<tr><td>160 Gd</td><td colspan="4">CD39</td></tr>
<tr><td>161 Dy</td><td colspan="4">NKG2A</td></tr>
<tr><td>162 Dy</td><td colspan="4">NKG2C</td></tr>
<tr><td>163 Dy</td><td colspan="4">EOMES</td></tr>
<tr><td>164 Dy</td><td colspan="4">CD28</td></tr>
<tr><td>165 Ho</td><td colspan="4">NKp30</td></tr>
<tr><td>166 Er</td><td>Fas-L</td><td>IgG4</td><td>Fas-L</td><td>IgG4</td></tr>
<tr><td>167 Er</td><td colspan="3">KIR2DL2/L3</td><td>CD103</td></tr>
<tr><td>168 Er</td><td colspan="3">TRAIL</td><td>KI67</td></tr>
<tr><td>169 Tm</td><td colspan="4">Tbet</td></tr>
<tr><td>170 Er</td><td colspan="4">CD3</td></tr>
<tr><td>171 Yb</td><td></td><td colspan="3">XCL1</td></tr>
<tr><td>172 Yb</td><td colspan="4">Perforin</td></tr>
<tr><td>173 Yb</td><td colspan="4">Granzyme B</td></tr>
<tr><td>174 Yb</td><td colspan="4">TOX</td></tr>
<tr><td>175 Lu</td><td colspan="4">TIGIT</td></tr>
<tr><td>176 Yb</td><td colspan="4">DNAM1</td></tr>
<tr><td>209 Bi</td><td colspan="4">CD16</td></tr>
</table>

Figure S2. NKG2A-expressing CD8 T cells are differentiated and possess TCR-independent NK-like functions in healthy donors. Related to Figure 3.

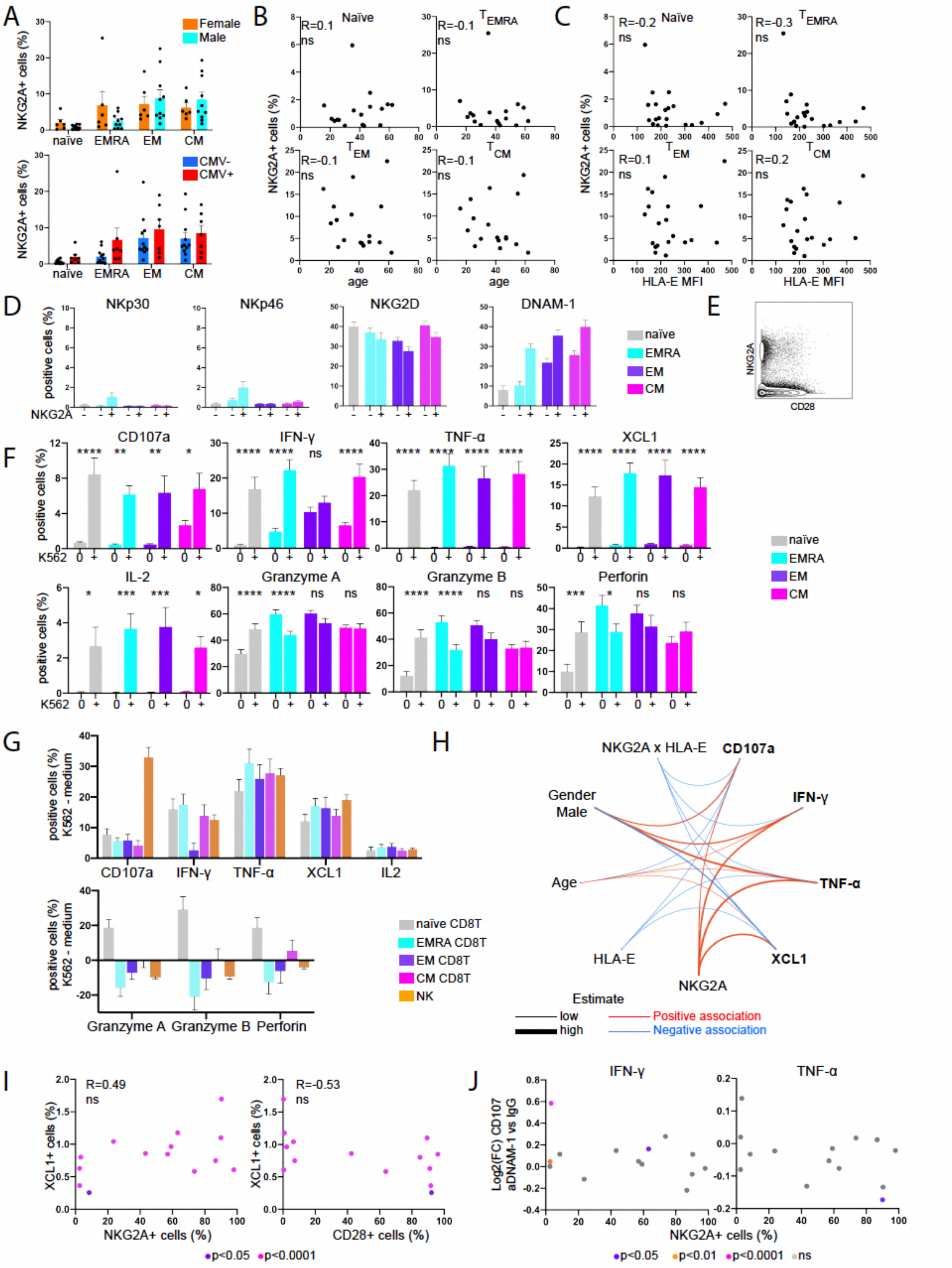

Figure S3. NKG2A defines an alternatively exhausted subset of TRM CD8+ T cells that retain TCR-independent functions in bladder tumors. Related to Figure 4.

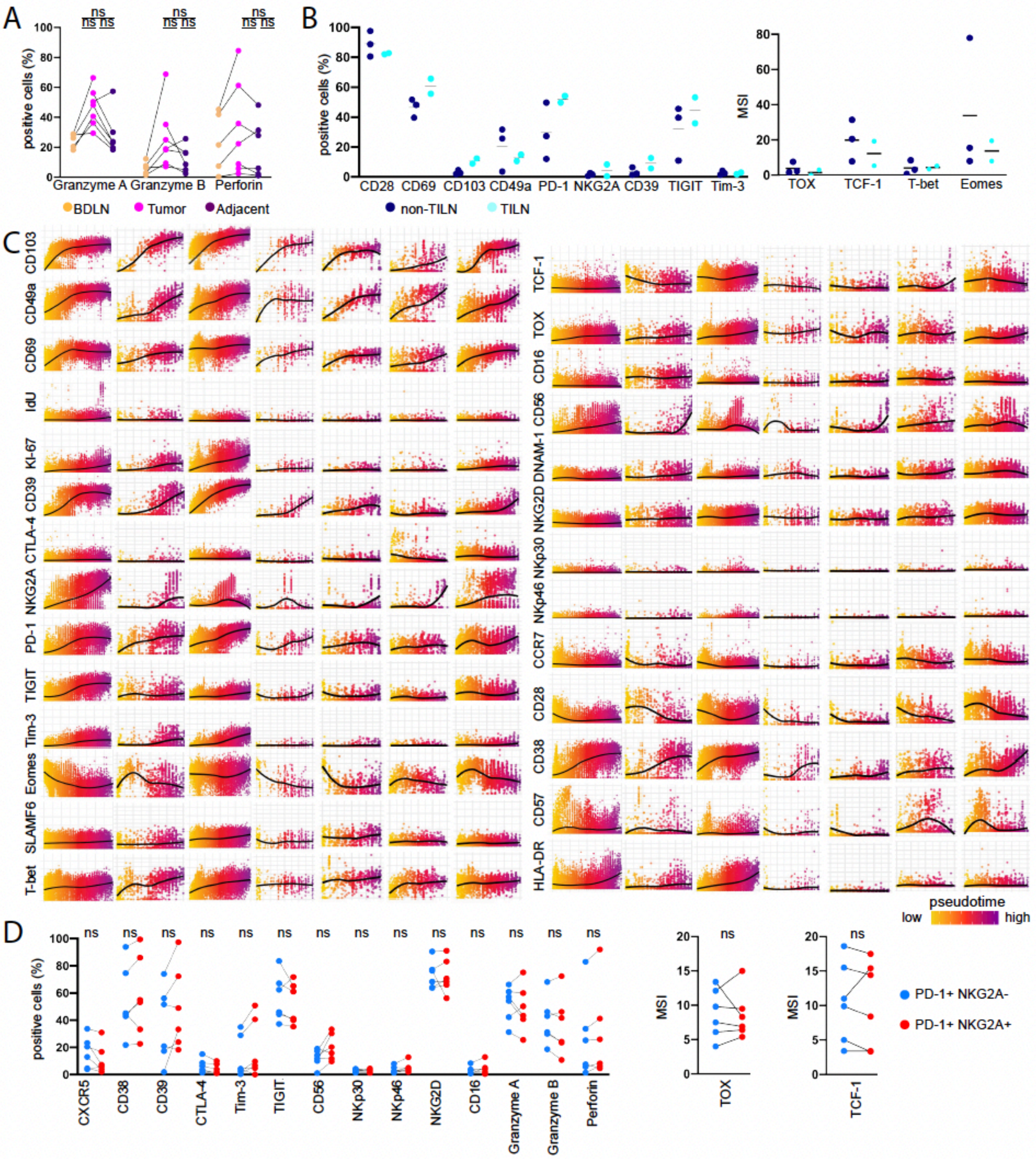

Figure S4. *KLRC1* (NKG2A) expression in CD8 T cells does not associate with a skewed clonal repertoire. Related to Figure 4.

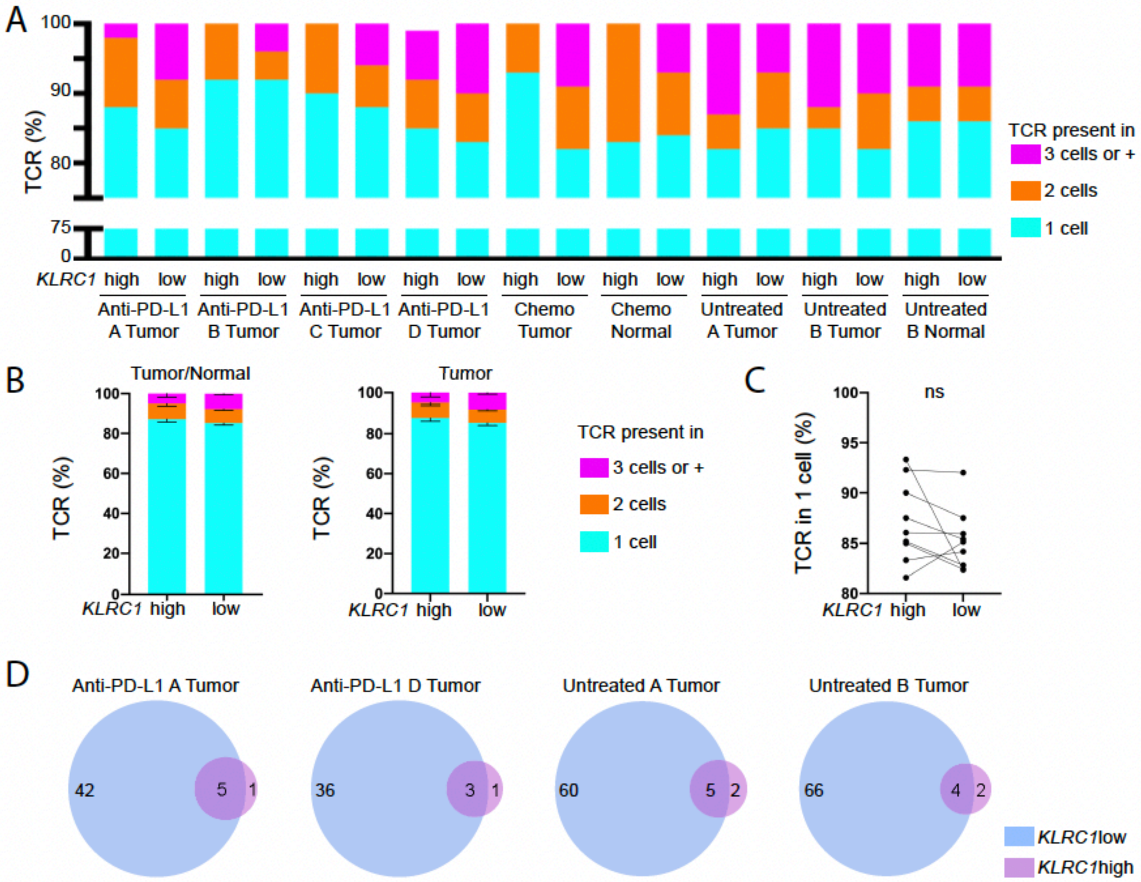

Figure S5. Compositional makeup of CD8 T cell clusters in bladder tumors according to pathology stage. Related to Figure 5.

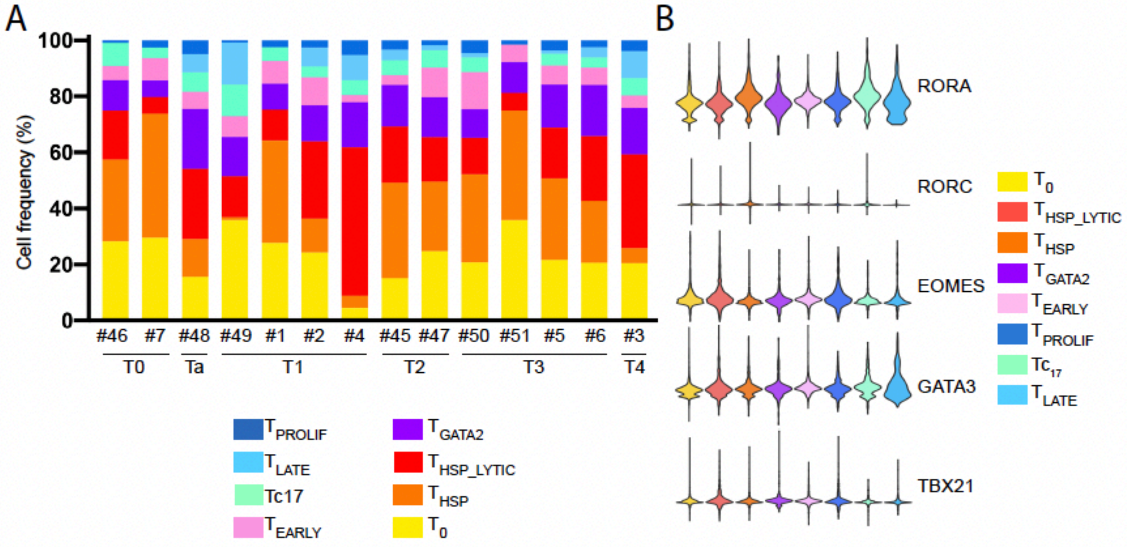

Figure S6. TGF-ß1 induces NKG2A expression on CD8 T cells and enhances their TCR-independent NK-like anti-tumor activity in response to NKG2A-blockade in bladder cancer. Related to Figure 6.

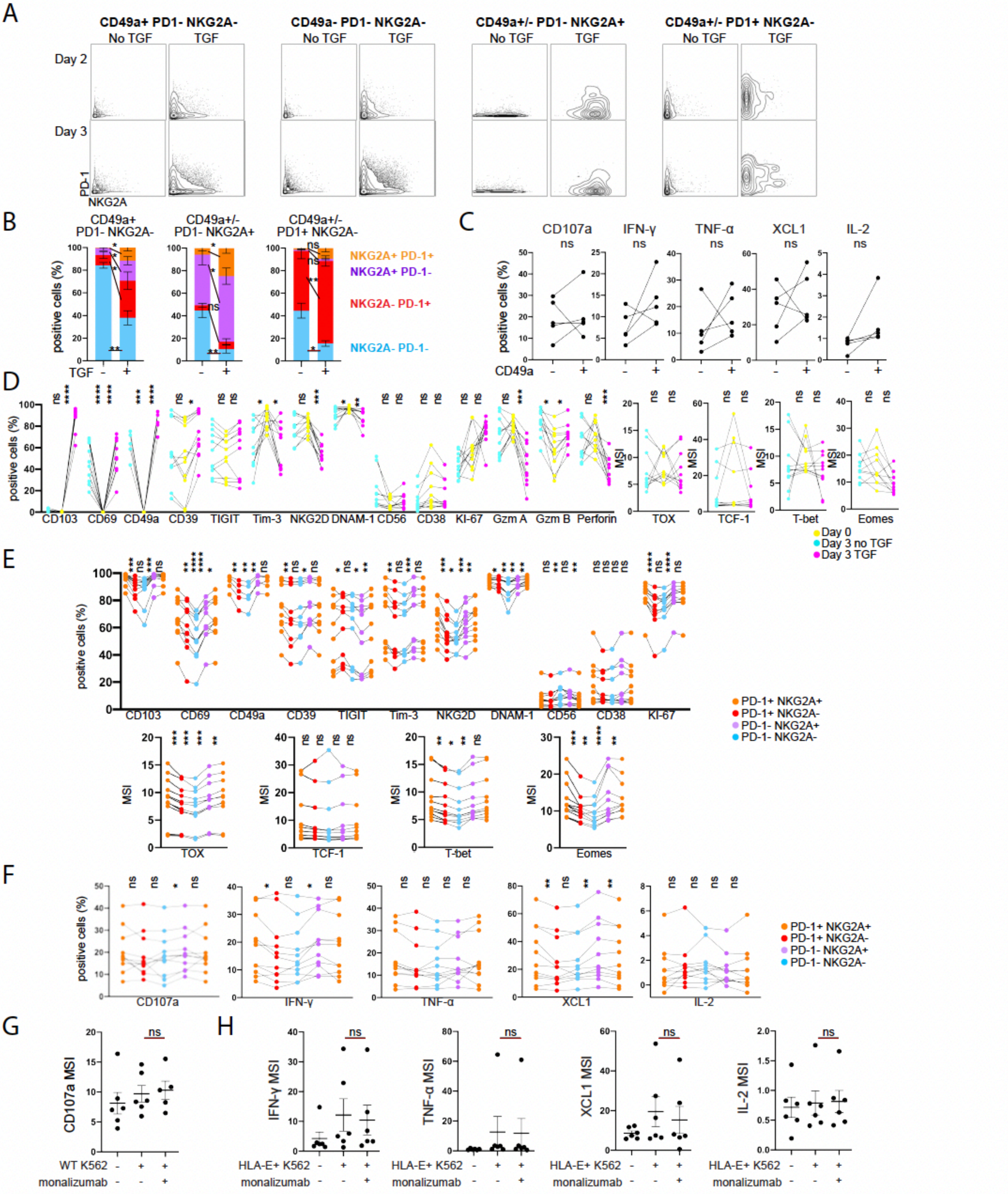
